# Repeat length increases disease penetrance and severity in *C9orf72* ALS/FTD BAC transgenic mice

**DOI:** 10.1101/2020.05.21.107425

**Authors:** Amrutha Pattamatta, Lien Nguyen, Hailey Olafson, Marina Scotti, Lauren A. Laboissonniere, Jared Richardson, J. Andrew Berglund, Tao Zu, Eric.T. Wang, Laura P.W. Ranum

**Author notes:** Corresponding author **Contact information of corresponding author** Laura P.W. Ranum, Ph.D., University of Florida, 2033 Mowry Road, Gainesville, FL 32610.

## Abstract

*C9orf72* ALS/FTD patients show remarkable clinical heterogeneity, but the complex biology of the repeat expansion mutation has limited our understanding of the disease. BAC transgenic mice were used to better understand the molecular mechanisms and repeat length effects of C9orf72 ALS/FTD. Genetic analyses of these mice demonstrate that the BAC transgene and not integration site effects cause ALS/FTD phenotypes. Transcriptomic changes in cell proliferation, inflammation and neuronal pathways are found late in disease and alternative splicing changes provide early molecular markers that worsen with disease progression. Isogenic sublines of mice with 800, 500 or 50 G_4_C_2_ repeats generated from the single-copy C9-500 line show longer repeats result in earlier onset, increased disease penetrance, and increased levels of RNA foci and RAN aggregates. These data demonstrate G_4_C_2_ repeat length is an important driver of disease and identify alternative splicing changes as early biomarkers of *C9orf72* ALS/FTD.

## Introduction

The intronic *C9orf72* G4C2 expansion mutation is the most common known genetic cause of both amyotrophic lateral sclerosis (ALS) and frontotemporal dementia (FTD) (DeJesus-Hernandez, Mackenzie et al., 2011, Renton, Majounie et al., 2011). Proposed molecular mechanisms include C9orf72 protein loss-of-function, RNA gain-of-function, and repeat associated non-ATG (RAN) protein toxicity (Ash, Bieniek et al., 2013, Chio, Restagno et al., 2012, DeJesus-Hernandez et al., 2011, Gijselinck, Van Langenhove et al., 2012, Mori, Weng et al., 2013, Nguyen, Cleary et al., 2019, Renton et al., 2011, Stewart, Rutherford et al., 2012, van der ZeeGijselinck et al., 2013, Van Langenhove, van der Zee et al., 2013, Van Mossevelde, van der Zee et al., 2017b, Zu, Liu et al., 2013). Numerous mouse models have been developed to better understand the relative contributions of loss-of-function and gain-of-function mechanisms in disease (Atanasio, Decman et al., 2016, Burberry, Suzuki et al., 2016, Chew, Gendron et al., 2015, Hao, Liu et al., 2019, Jiang, Zhu et al., 2016, Liu, Pattamatta et al., 2016, O’Rourke, Bogdanik et al., 2015, Peters, Cabrera et al., 2015, Schludi, Becker et al., 2017, Zhang, Gendron et al., 2018, Zhang, Guo et al., 2019). While C9orf72 protein levels measured in autopsy samples are lower in C9 patients (Waite, Baumer et al., 2014) *C9orf72* knockout mice develop peripheral immune phenotypes but not ALS/FTD related phenotypes (Atanasio et al., 2016, Burberry et al., 2016, Jiang et al., 2016, O’Rourke et al., 2015) making it unlikely that C9orf72 loss of function alone is a major driver of disease. In contrast, *Drosophila* and mouse models that overexpress specific RAN proteins develop neurodegenerative and motor phenotypes (Atanasio et al., 2016, Burberry et al., 2016, Chew et al., 2015, Hao et al., 2019, Jiang et al., 2016, Liu et al., 2016, Mizielinska, Gronke et al., 2014, O’Rourke et al., 2015, Peters et al., 2015, Schludi et al., 2017, Zhang et al., 2018, Zhang et al., 2019) indicating RAN proteins can be toxic and may play a role in disease. RNA gain-of-function effects may cause RNA processing abnormalities that contribute to disease through the sequestration of RNA binding proteins by the repeat-containing sense and antisense transcripts. Studies on C9 iPSC derived neurons (iPSNs) and C9-ALS autopsy tissue have reported transcriptomic changes (Donnelly, Zhang et al., 2013, Ebbert, Ross et al., 2017, Prudencio, Belzil et al., 2015). Additionally, a number of RNA binding proteins (RNA-BPs) that interact with short stretches of G_4_C_2_ repeats have been identified through unbiased interactome screens, including Pur-α, ADARB2, hnRNPH, hnRNPA1, hnRNPA2/B1, ALYREF, nucleolin and RanGAP1 (Cooper-Knock, Walsh et al., 2014, Donnelly et al., 2013, Freibaum, Lu et al., 2015, Haeusler, Donnelly et al., 2014, Jovicic, Mertens et al., 2015, Lee, Chen et al., 2013, Sareen, O’Rourke et al., 2013, Xu, Poidevin et al., 2013, Zhang, Donnelly et al., 2015). Crosslinking immunoprecipitation (CLIP) analyses using autopsy material from the frontal cortex of C9-ALS patients shows that hnRNP-H binds to G_4_C_2_ transcripts with short repeats (Conlon, Lu et al., 2016). Since there is little consensus on which RNA binding proteins are sequestered by the G_4_C_2_ repeats, the role of RNA gain-of-function mechanisms in C9orf72 ALS/FTD remains unclear.

There is remarkable clinical heterogeneity among *C9orf72* expansion carriers with clinical presentations ranging from muscle wasting characteristic of ALS in some patients, to disinhibition and cognitive deficits characteristic of FTD in others. While some expansion carriers remain asymptomatic into their 90s, the frequency of reduced penetrance is not yet clear because patients with symptoms are much more likely to be seen by a physician and genetically tested compared to asymptomatic mutation carriers (Majounie, Renton et al., 2012, Murphy, Arthur et al., 2017). Repeat length and somatic repeat instability, which are known to contribute to Huntington disease and other repeat expansion disorders (Jones, Houlden et al., 2017, Lee, Wheeler et al., 2015, Wright, Collins et al., 2019), may contribute to the reduced disease penetrance of *C9orf72* ALS/FTD, differences in age-of-onset and the wide-ranging clinical effects of the *C9orf72* expansion mutation (Chio et al., 2012, Murphy et al., 2017). However, because of ascertainment bias, technical difficulties in measuring repeat length and somatic instability in patients, it is challenging to study the effects of repeat length as a modifier of *C9orf72* ALS/FTD (Hsiung, DeJesus-Hernandez et al., 2012, Van Mossevelde, van der Zee et al., 2017a).

To better understand the molecular mechanisms of disease, we and others generated bacterial artificial chromosome (BAC) transgenic mouse models that show molecular phenotypes of the disease including sense and antisense RNA foci and RAN protein aggregates, although the relative levels of these molecular phenotypes have not been directly compared (Jiang et al., 2016, Liu et al., 2016, O’Rourke et al., 2015, Peters et al., 2015). Three of these models, which were developed on B6 genetic backgrounds showed no (O’Rourke et al., 2015, Peters et al., 2015) or subtle hippocampal degeneration and social behavioral deficits (Jiang et al., 2016) but not typical features of ALS/FTD. In contrast, several independent lines of BAC transgenic mice generated on the FVB background (Liu et al., 2016) showed both the molecular and behavioral features of ALS/FTD including open field and DigiGait abnormalities, paralysis, motor neuron loss and decreased survival (Liu et al., 2016).

Here we describe the transgene integration sites of these C9-BAC lines (Liu et al., 2016), further establishing that the ALS/FTD phenotypes in these mice occur independent of integration effects. RNAseq analyses, using the most penetrant single copy C9-500 line, show that transcriptomic profiles are distinct at different stages of disease and that alternative splicing abnormalities are prevalent prior to onset of overt disease features suggesting their potential utility as early biomarkers of ALS/FTD. Using the single copy C9-500 line, we generated an allelic series of mice containing 800, 500 or 50 repeats and demonstrate that longer repeat tracts in an isogenic background increase disease penetrance and decrease age-of-onset and survival. Taken together, these data demonstrate that ALS/FTD phenotypes in FVB C9-BAC mice are driven by gain-of-function effects of the expansion mutation, these effects occur independent of integration site and that repeat length is a major driver of disease.

## Results

### Phenotypes in C9-BAC mice independent of integration sites

We previously reported the development of a BAC transgenic model of C9orf72 ALS/FTD on the FVB/NJ background (Liu et al., 2016). Four independent lines were generated by pronuclear injection of a circularized BAC containing a 98.3 kb human DNA insert, including a large G_4_C_2_ repeat-expansion mutation (Fig 1A). To further characterize these mice, we performed whole genome sequencing to determine transgene break points, genomic integration sites, and number of transgene copies for each of the four BAC mouse lines. Transgene break and genomic integration sites were identified by computational analyses of discordant read-pairs from transgenic DNA compared to mouse and human reference genomes. Transgene copy number for each of the lines was determined by comparing the regional coverage depth of the BAC with the average coverage depth of the mouse reference genome. These data show that the transgenes in all four BAC lines were inserted into distinct single integration sites. Transgene copy number for each of the lines was consistent with previous estimates based on Southern blot and qRT-PCR analyses (Liu et al., 2016).

**Figure 1:**
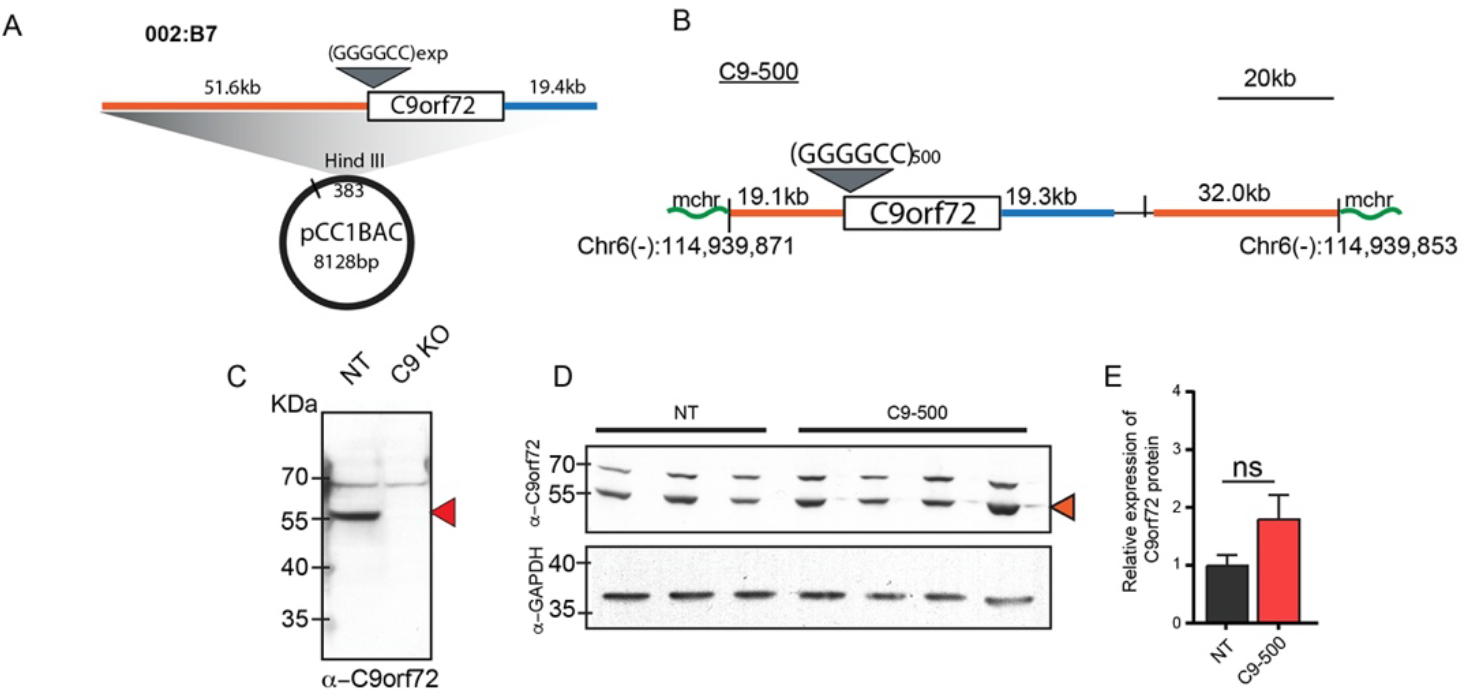
Characterization of transgene integration site, RAN protein expression and C9orf72 protein levels in C9-BAC mice. A) BAC construct containing human C9orf72 gene and 51.6 kb and 19.4 kb of upstream (orange) and downstream (blue) flanking sequence, respectively. B) Map showing breakpoint and integration of the BAC in an intergenic region on mouse chromosome 6. C, D) Protein blots probed with α-C9ORF72 antibody. E) Protein blot quantitation. Data information: Statistical analyses for panel E were performed using one-way ANOVA with Bonferroni correction for multiple comparisons with mean +/− SEM shown; not significant (ns) = p > 0.05. See also Figure S1, 2 and 3.

In the single-copy C9-500 line, the BAC transgene inserted between Chr 6(−): 114,939,871 and Chr 6(−): 114,939,853 in the mouse genome resulting in an ~18 bp deletion in an intergenic region ~18.1 kb distal and ~64.5 kb proximal of the nearest flanking genes *Vgll4* and *Tamm41*, respectively (Fig 1B). The breakpoint on the circular BAC transgene occurred 19,097 bp upstream of *C9orf72*. At the site of integration, the C9-500 line contains the full length *C9orf72* gene containing ~500 G_4_C_2_•G_2_C_4_ repeats with ~19.1 kb of 5’ human flanking sequence and ~19.4 kb of 3’ human flanking sequence. The remaining sequence includes the pCC1BAC backbone followed by 32.0 kb of human DNA originally 5’ of the breakpoint on the BAC. In summary, the BAC transgene in the C9-500 line has a single integration site containing a full-length copy of the *C9orf72* gene, substantial human flanking sequence and ~500 G_4_C_2_•G_2_C_4_ repeats.

In the C9-500/32 line, two copies of the transgene were integrated between mouse chromosome 18 (−) 17,919,900 and, and the flipped chromosome 18 (+) strand at position 18,526,504 (Fig S1A). No annotated genes were found in this region and no decrease in coverage between the breakpoints was observed indicating a small mouse chromosomal rearrangement but not a mouse chromosomal deletion occurred in this region (Fig S1A). In the C9-500/32 line, the first copy of the integrated human transgene contains 46.6 kb of upstream and 19.4 kb of downstream flanking sequences and the full-length copy of the *C9orf72* gene. The second transgene copy contains the full 51.6 kb upstream flanking region and a small portion of *C9orf72* which terminates 3’ of the repeat within the first intron. Southern blot analyses show that the larger repeat expansion is located in the full length *C9orf72* copy and the shorter expansion in the second copy containing the truncated *C9orf72* gene (Fig S1B, C).

The C9-36/29 line has three copies of the full-length *C9orf72* gene, and one truncated copy inserted into the second intron of *metallophosphoesterase 1 (Mppe1)* (Fig S2A). Although the deletion of exons 3-4 of *Mppe1* is predicted to lead to the expression of a truncated protein lacking the N-terminal region and a portion of the metallophosphoesterase domain, qRT-PCR and RNA sequencing detected no significant differences in transcript levels including over exons 3 and 4 (Fig S2B, C). While it is possible that a truncated metallophosphoesterase protein is expressed and could cause some type of deleterious effect, no overt phenotypic differences were found in this line compared to the other phenotypic lines.

The C9-37 line has a single transgene insertion site on chromosome 4 at a position with no annotated genes. This insertion contains a partial copy of *C9orf72* extending from exon 1 into intron 9 plus 19.5 kb of endogenous human upstream flanking sequence. Due to the position of the transgene break, the pCC1BAC backbone plus an additional 17.3 kb of 3’ sequence is integrated further upstream of *C9orf72* (Fig S2D). The C9-37 line, which lacks the 3’ end of *C9orf72*, is the only line that does not contain a full-length copy of the transgene, and the only line that does not develop overt ALS/FTD phenotypes.

We measured the levels of exon1a containing sense RNA transcripts by RT-qPCR and polyGP RAN protein using a meso scale discovery (MSD) assay (Fig S3). RT-qPCR shows that the levels of sense expansion containing RNA transcripts in the brain from each of four C9-BAC lines correlate with transgene copies (Fig S3A). In contrast, the levels of polyGP RAN proteins are the highest in the most penetrant C9-500 line in both the cortex and cerebellum (Fig S3B).

In summary, three independent *C9orf72* BAC lines show similar ALS/FTD phenotypes (Liu et al., 2016), strongly supporting the hypothesis that these phenotypes are caused by the repeat expansion mutation and not disruptions of integration site genes or other changes at the various integration sites. Of the two lines with relatively short repeats (C9-36/29 vs C9-37), only the C9-36/29 line containing 4 copies of transgenes develops ALS/FTD phenotypes. This difference could result from the higher expression of the relatively small G_4_C_2_ expansions in the phenotypic C9-36/29 line (4 transgene copies) compared with the C9-37 line (1 transgene copy) or the truncation of the *C9orf72* gene found in the C9-37 line.

### C9orf72 protein overexpression not associated with disease in C9-500 mice

To study if the integration of the transgene results in an upregulation of C9orf72 protein levels that might contribute to phenotypes seen in the C9-500 line, we compared the levels of the C9orf72 protein in 20-week-old C9-500 mice with age-matched NT mice. Splice variants of the *C9orf72* mRNA generate two isoforms of the protein with predicted sizes of ~55 kDa and ~35 kDa (Xiao, MacNair et al., 2015). Commercially available antibodies detect the long isoform. We used the C9orf72 Genetex antibody that detects both human and mouse C9orf72 proteins (Laflamme, McKeever et al., 2019) and confirmed antibody specificity by showing that the 55kDa protein is detected in the control but not *C9orf72* KO brain lysates (generously donated by Dr. Robert. H. Baloh) (Fig 1C). While the levels of the C9orf72 protein trended towards an increase in C9-500 mice, no significant upregulation was detected in cortical brain lysates from C9-500 compared to NT mice (Fig 1D, E).

In summary, the phenotypic C9-500 mice show modest but insignificant elevation of C9orf72 protein suggesting that this change is unlikely to be a critical driver of the ALS/FTD phenotypes in this mouse model.

### Neuroinflammatory transcriptome changes predominate at end-stage

RNA dysregulation is thought to play a role in *C9orf72* ALS/FTD (Butti & Patten, 2018, Prudencio et al., 2015, Prudencio, Gonzales et al., 2017) but little is known about how the transcriptome is affected early in the course of disease or how these changes progress over time. To look for transcriptomic abnormalities that occur during disease progression, we performed RNA sequencing on frontal cortex samples from ten female C9-500 mice at 20 weeks of age with no overt cage behavior abnormalities (C9+ presymptomatic), four C9+ animals that developed acute rapidly progressive phenotypes (20-22 weeks old) (Acute) and three NT controls.

Sample to sample correlation based on differential gene expression measured using DESeq2 shows that the acute cohort of C9-500 mice have unique gene expression profiles. Additionally, global gene expression changes in the NT animals are similar to the C9(+) animals but acute animals were significantly different from both the NT and C9(+) presymptomatic animals (Fig 2A). In the Acute mice, we identified 2514 upregulated and 2921 downregulated genes compared to NT animals, with false discovery rate (FDR) <0.05 (Fig 2A). Heat map of the top 50 significant genes in the Acute vs NT mice are shown in Fig S4. Gene ontology (GO) analyses in the Acute vs NT mice shows that the upregulated pathways include negative regulation of cell proliferation, inflammatory response and actin cytoskeleton organization, (p values indicated on the left) (Fig 2B). Significantly downregulated pathways include brain development, synaptic organization and neuron projection development (Fig 2B). Since the Acute C9-BAC mice mimic the neuropathology seen in end-stage C9-ALS patients, we compared the gene expression profiles of the cortex from Acute mice and C9-ALS patients. RNA sequencing data obtained from the Prudencio et al. study (Prudencio et al., 2015) was reanalyzed using STAR (Dobin, Davis et al., 2013) for alignment Kallisto (Bray, Pimentel et al., 2016) to obtain transcript per million values. Using these parameters, we identified 36 genes that were consistently dysregulated between C9-ALS and unaffected individuals in this dataset. Of these, 15 of the 36 were also dysregulated in our acute mice, including SerpinH1 (Fig S5). This gene belongs to the serine protease inhibitor family, and several members of this gene family, including SerpinA3 and SerpinA1 are hypothesized to disrupt neuronal function and have been found to be differentially expressed in C9orf72 ALS patient autopsy tissue (Ebbert et al., 2017, Prudencio et al., 2015).

**Figure 2:**
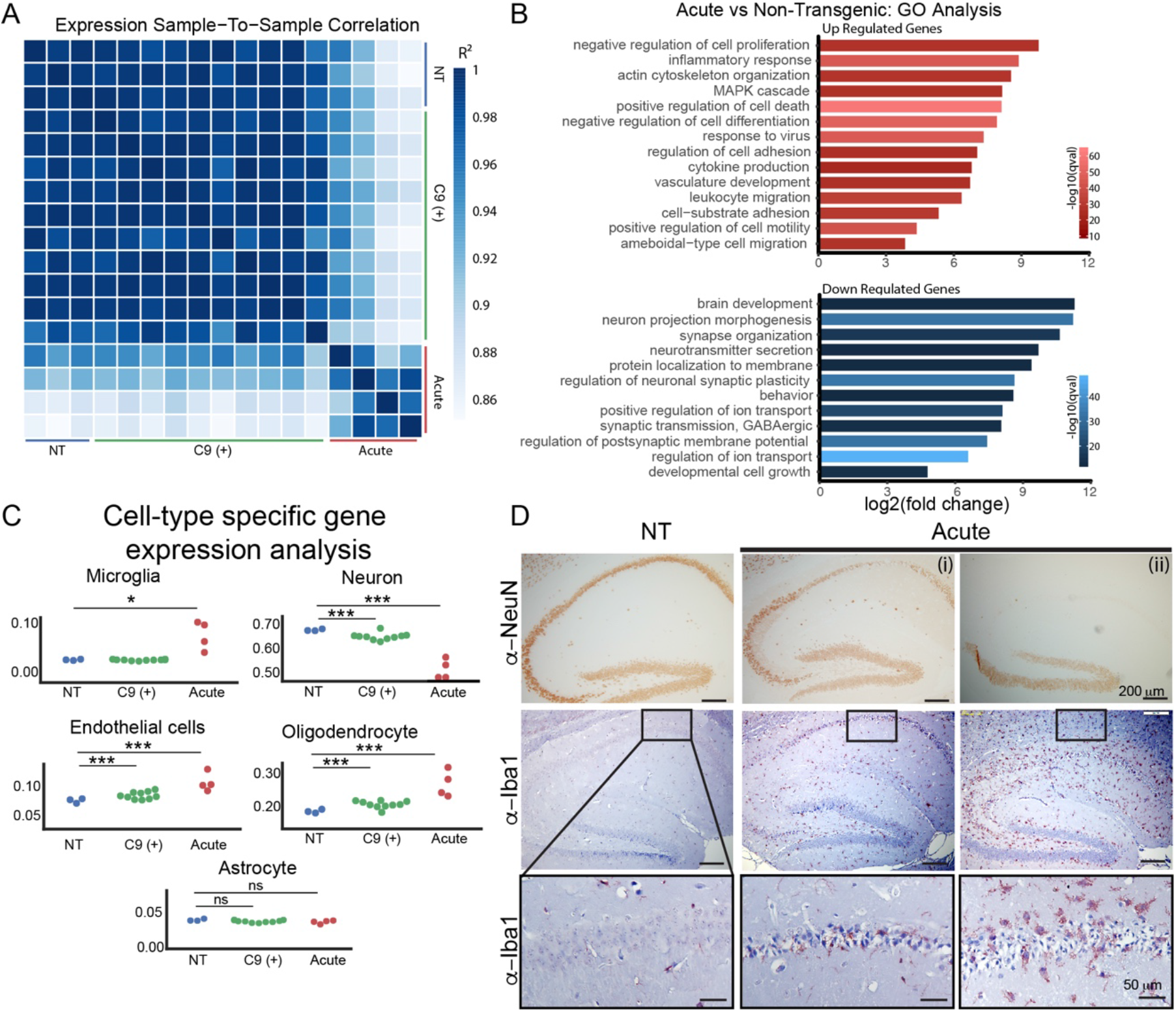
Prominent neurodegenerative and neuroinflammatory changes in acute end-stage C9-BAC mice. A) Sample-to-sample correlation plot shows increased correlation in gene expression changes among Acute mice and C9(+) presymptomatic mice but not between groups. B) Gene ontology analyses of gene expression changes in Acute vs NT mice. C) Cell type enrichment analyses in NT, C9(+) presymptomatic and Acute mice. Statistical tests and significance are shown in Table 1. D) Immunohistochemistry of Acute C9-BAC mice stained for neuronal (NeuN) and microglial (Iba1) markers. Inset shows zoom-in of microglial staining. Acute (i) and (ii) represent earlier and later disease stages in C9-BAC mice, respectively. See also Figure S4,5,6.

Since a large number of neuroinflammatory and neurodegenerative gene expression changes in the Acute cohort may have resulted from inflammatory and neurodegenerative processes that cause loss of neurons or increases in inflammatory cells, we estimated cell type changes in each sample. For this analyses, we used publicly available datasets that had characterized gene expression in seven different cell types (neurons, microglia, astrocytes, endothelial cells, oligodendrocyte precursor cells, myelinating oligodendrocytes and newly formed oligodendrocytes) in the mouse brain (Zhang, Chen et al., 2014). Because there is a considerable overlap between the expression profiles of oligodendrocyte precursor cells, myelinating oligodendrocytes and newly formed oligodendrocytes, we combined these cell types into a single category called “oligodendrocytes”. We observed an increase in the estimated proportion of microglial, oligodendrocyte and endothelial cells in acute vs NT mice, and a decrease in estimated proportion of neurons in Acute vs NT mice (Fig 2C, Table 1). Additionally, significant albeit small differences were also observed between C9(+) presymptomatic and NT mice in the estimated proportion of endothelial, oligodendrocyte and neuronal cells, but not microglia (Fig 2C, Table 1). Consistent with these findings, IHC shows overt loss of immunoreactivity to the neuronal marker NeuN and increased staining of the microglial marker Iba1 in the acute mice compared to NT mice (Fig 2D). Interestingly, cresyl violet staining of C9(+) presymptomatic animals showed no overt pathology in the hippocampus (Fig S6). No significant differences were observed in the estimated proportion of astrocytes in acute or C9(+) mice relative to NT (Fig 2C).

**Table 1:**
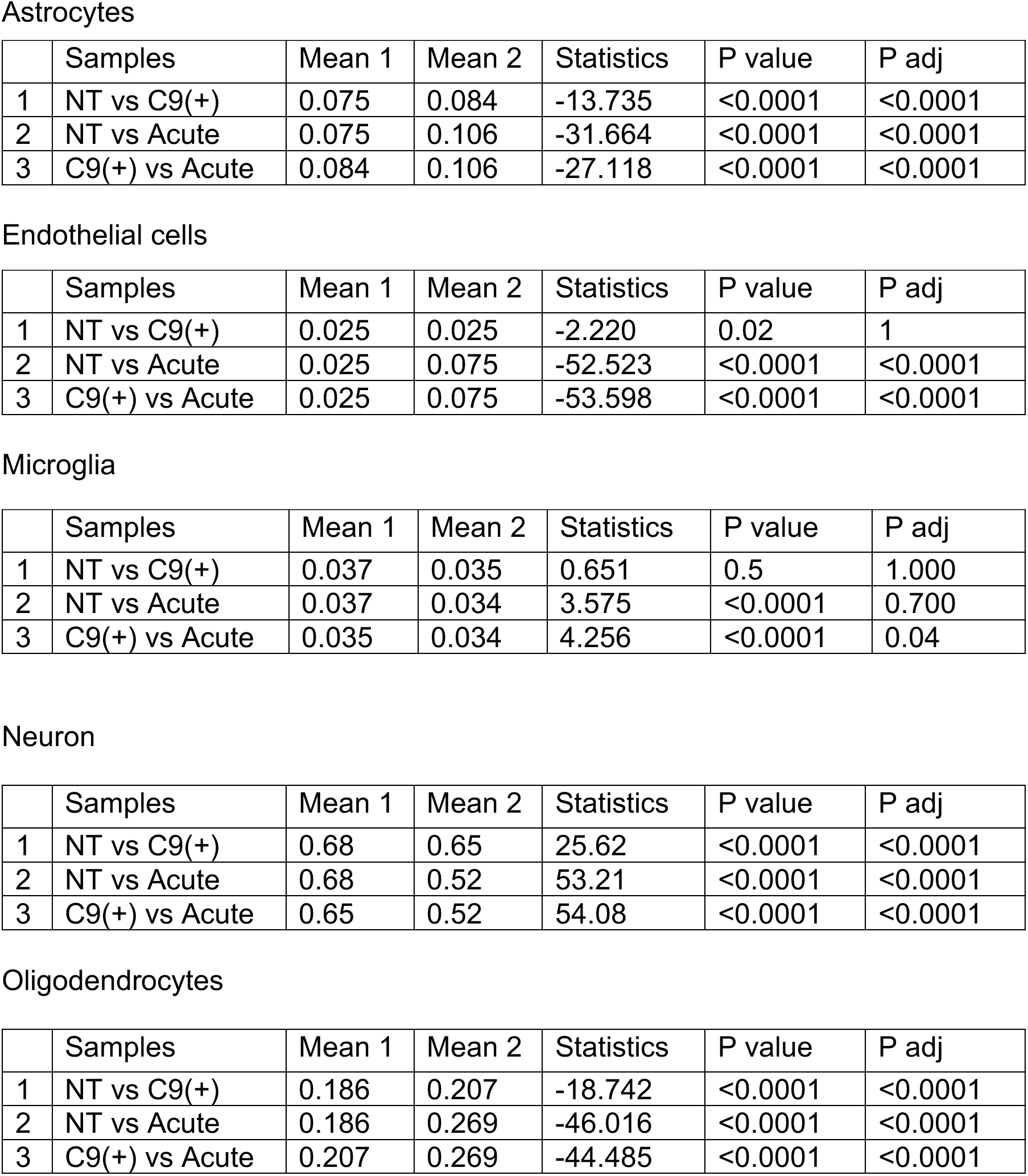
Cell type analysis on transcriptome data from NT, C9(+) presymptomatic and Acute mice.

Taken together, robust changes in gene expression were seen in Acute mice compared to NT littermates. These changes are consistent with the neuronal loss and increased numbers of microglia in Acute compared to NT mice. Similarly, presymptomatic C9(+) animals showed modest changes in cell type specific genes associated with neurons, endothelial cells and oligodendrocytes compared to NT mice.

### Alternative Splicing Changes Characterize Disease States in ALS/FTD

Changes in alternative splicing and alternative polyadenylation were previously reported in autopsy brains from C9-positive ALS but not C9-negative sporadic ALS cases (Prudencio et al., 2015). We examined alternative splicing variants in our mouse model using MISO analyses to identify changes that occur during disease progression. Although very few differentially expressed genes were detected in presymptomatic animals, 240 and 539 genes showed alternatively splicing changes in C9(+) presymptomatic vs NT and C9(+) Acute vs NT cohorts, respectively (Fig 3A). These results demonstrate that alternative splicing abnormalities are an early molecular signature of C9orf72 ALS/FTD. Eighty-three of these alternative splicing events were shared between the Acute and C9(+) presymptomatic cohorts (Fig 3A) and the percent spliced in (psi) values of these shared events increase with disease progression (Fig 3B). Elavl2, an RNA binding protein enriched in the neurons that affect neuronal excitability (Ince-Dunn, Okano et al., 2012), was found to be alternatively spliced in both C9(+) presymptomatic animals and Acute animals and the psi value increased with disease progression (Fig 3C). A summary of the alternatively spliced events found in both C9(+) presymptomatic and Acute animals is shown in Fig 3D. Because the psi values of these 83 genes are also changed in the presymptomatic mice and the psi values continue to increase with disease progression. Although beyond the scope of this study, these genes may be useful as biomarkers to monitor disease progression in C9-ALS/FTD patients (Fig 3D, Table 2).

**Table 2:**
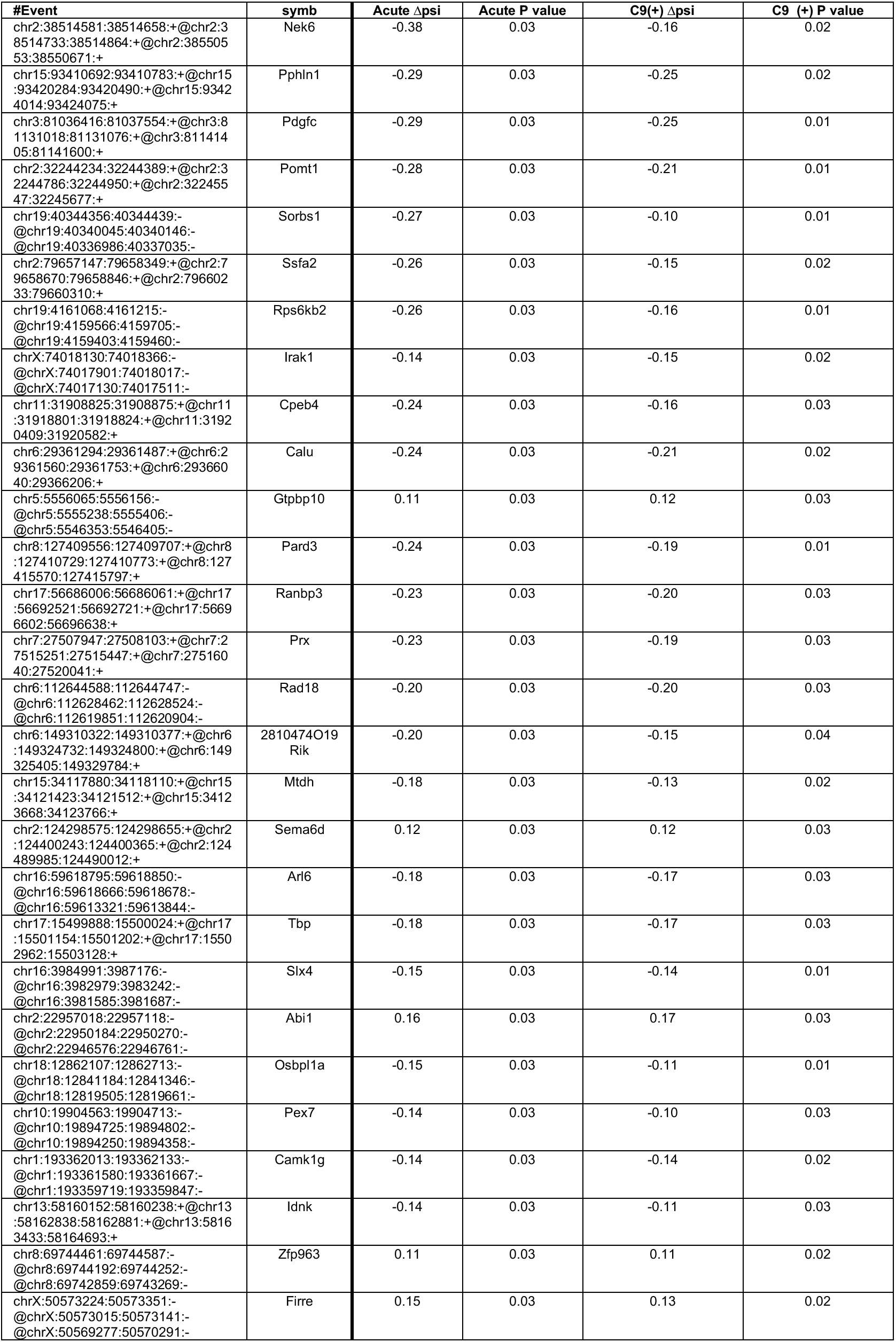
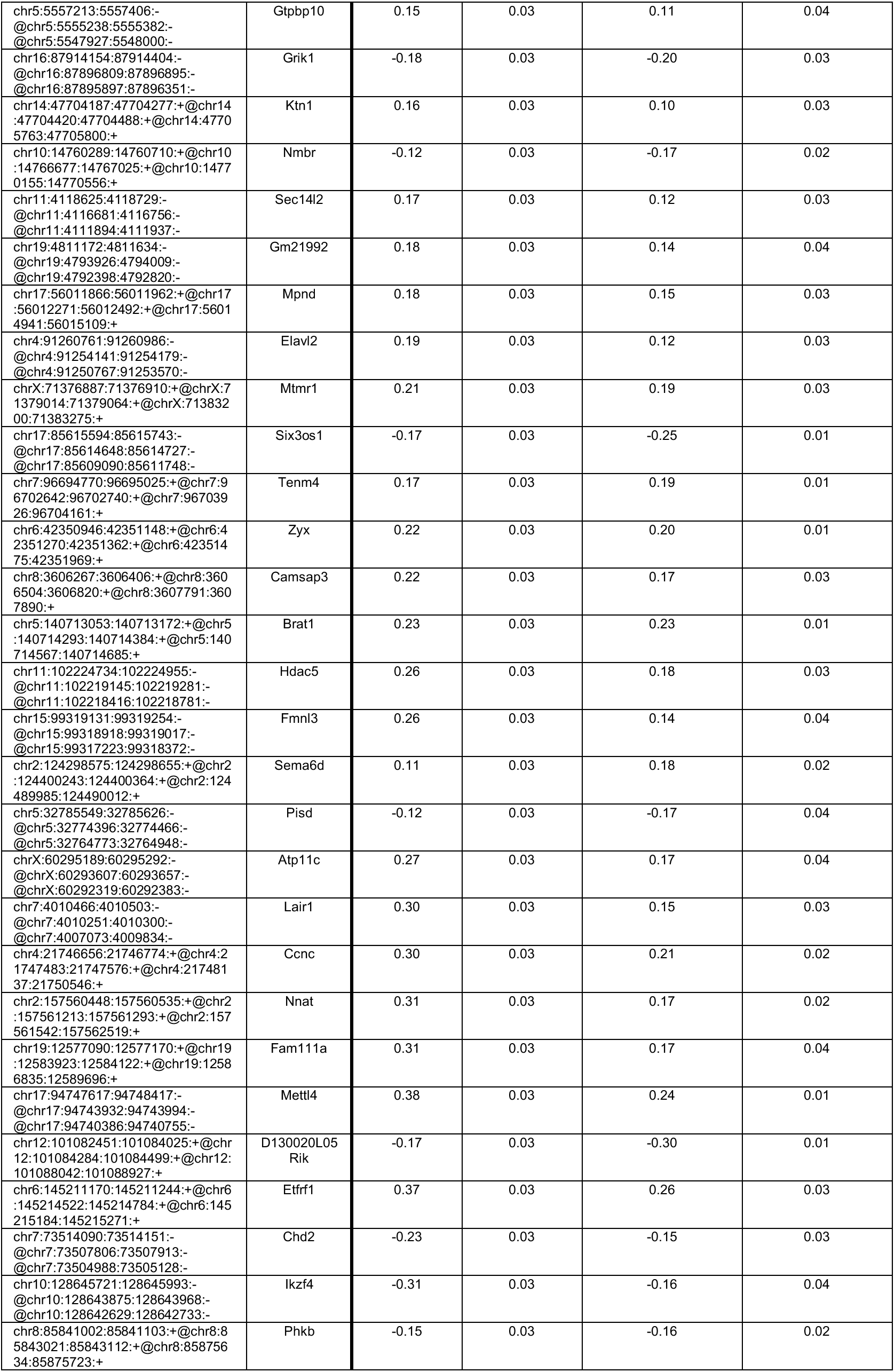
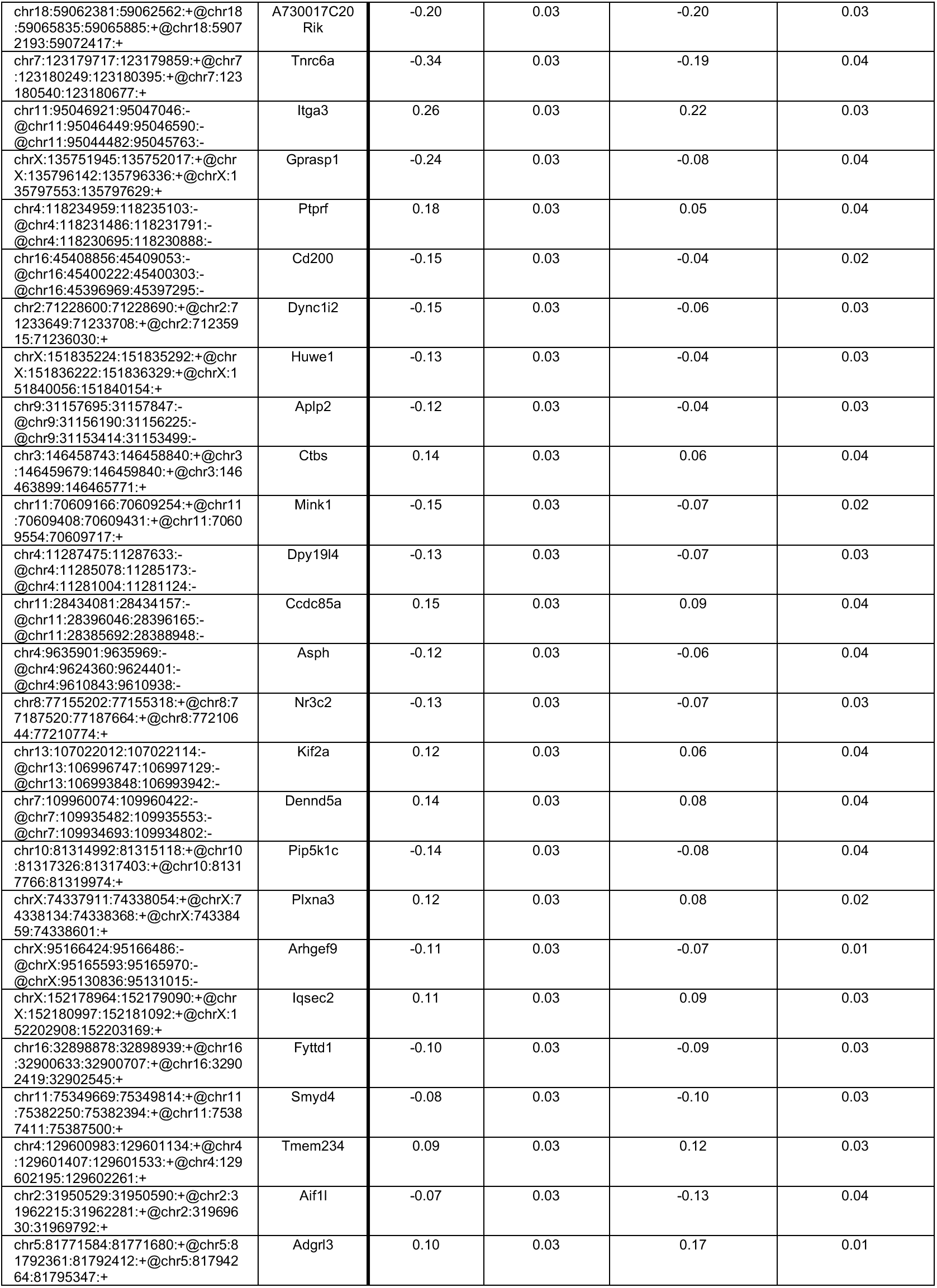
Common alternative splicing events in C9(+) presymptomatic and Acute mice.

**Figure 3:**
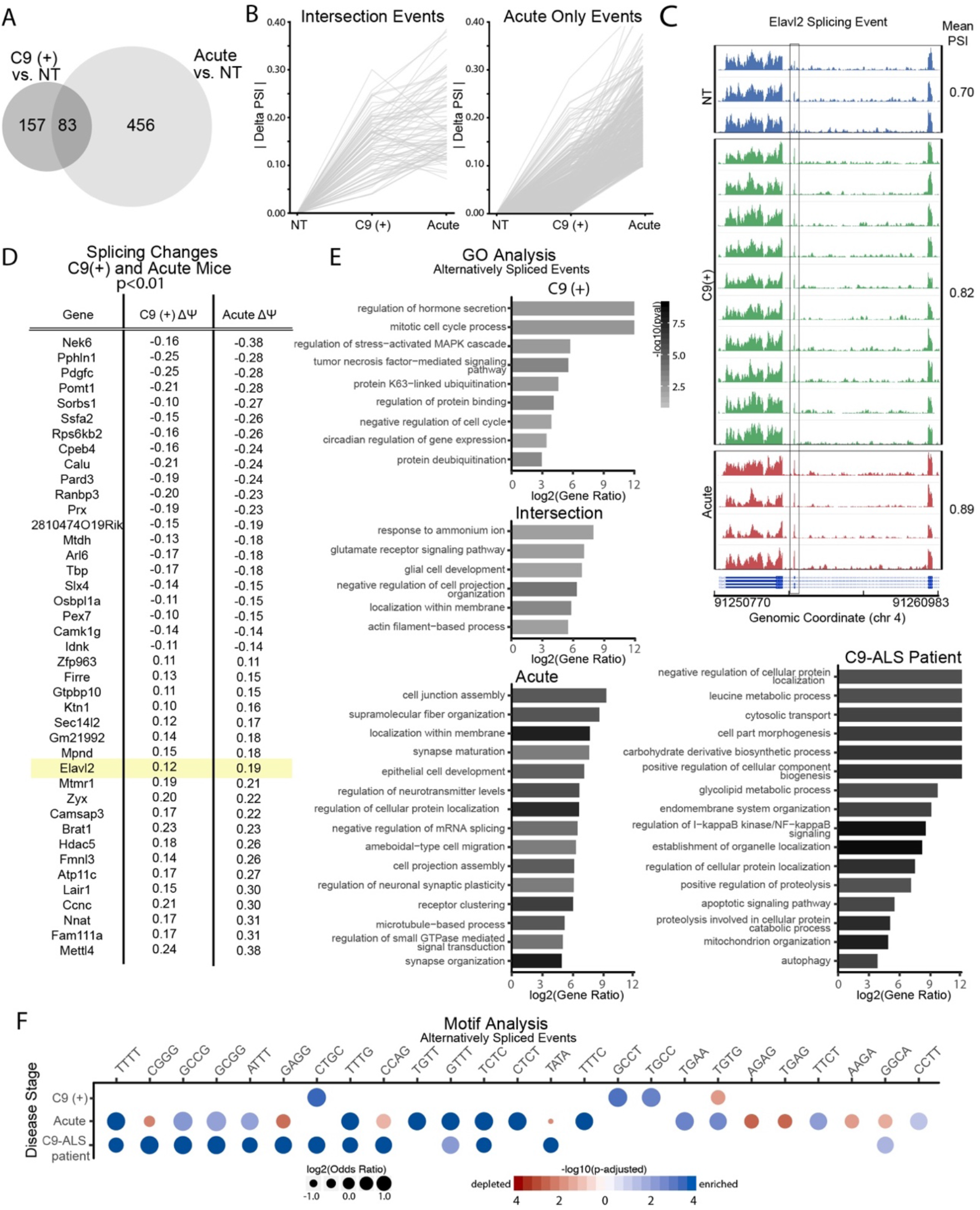
Abundant alternative splicing changes in C9-BAC mice. A) Venn diagram showing number of alternative splicing changes in Acute and presymptomatic mice compared to NT mice. B) Alternative splicing events found in both presymptomatic and Acute animals show delta psi values for a group of markers increase with disease severity. C) *Elavl2* is alternatively spliced in both the C9(+) presymptomatic [C9(+)] and Acute mice. D) Gene IDs and corresponding Δpsi values in C9(+) presymptomatic and Acute animals. Additional information on alternative splicing events is provided in Table 2. E) Gene Ontology categories are shown for alternatively spliced events found only in: C9(+) presymptomatic mice, both presymptomatic and Acute animals (intersection), only Acute animals and C9-ALS patients. F) Motif analysis of alternative splicing events in presymptomatic C9(+) mice, Acute C9(+) mice and C9-ALS patients shows enrichment of motifs. See also Figure S7.

GO analyses were used to better understand the categories of alternatively spliced genes that show changes in presymptomatic and Acute mice and changes common to presymptomatic and acutely affected animals (Fig 3E). MISO was also performed to detect alternative splicing changes in previously published RNAseq data from human autopsy tissue (Fig 3E). Splicing changes occurring in both acutely affected mice and end-stage C9-ALS patients were enriched for several similar GO categories including genes with alternative splicing abnormalities that are normally involved in neuronal death, oxidative stress, cytoskeletal pathways and inflammation. In contrast, pathways dysregulated in presymptomatic mice include synaptic transmission and membrane localization. In summary, we show that alternative splicing perturbations occur early in disease and progressively worsen with increases in disease severity.

### Motif analyses of intronic regions flanking exons dysregulated in C9 ALS/FTD

In myotonic dystrophy, a well-established RNA gain-of-function disease, CUG or CCUG expansion RNAs sequester MBNL proteins into intranuclear foci, thus preventing their normal function in regulating posttranscriptional processing including alternative splicing and polyadenylation (Scotti & Swanson, 2016). As expected, sequence analyses of abnormally spliced exons from DM1 skeletal muscle are most significantly enriched for MBNL YGCY binding motifs (Fig S7).

In contrast to DM1, sequence analyses of abnormal splicing events in C9(+) presymptomatic mice and acute mice shows the enrichment of a larger number and more diverse set of tetramer motifs, including AT and multiple types of GC rich repeat motifs (Fig 3E). A diverse set of repeat motifs are also enriched in the C9-ALS patient splicing data (Fig 3E). In contrast to the prominent involvement of a single category of RBP in DM1, the multiple types of RNA binding motifs found in the dysregulated genes in C9 ALS/FTD are consistent with published literature and indicate that C9orf72 RNA dysregulation may involve a larger group of RNA binding proteins, or may reflect concurrent processes that prevent unambiguous identification of a single RBP driver of disease (Fig 3E).

In summary, we show that alternative splicing changes are found in C9-BAC mice and increase with disease severity and hence may provide useful biomarkers and tools to understand disease progression. In contrast to DM1, motif enrichment analyses shows that the alternative splicing changes found in our C9-BAC mice are unlikely to be caused by sequestration of a single category of RBP.

### Repeat length increases penetrance and decreases survival in allelic series of C9-BAC mice

Repeat length is a known modifier of disease severity in multiple repeat expansion diseases including Huntington disease, DM1 and multiple spinocerebellar ataxias (Gusella & MacDonald, 2000, Paulson, 2018). Somatic instability and technical difficulties measuring G_4_C_2_ repeat length in human *C9orf72* patients have made the contribution of repeat length to age of onset and disease severity in *C9orf72* ALS/FTD difficult to determine. To test the hypothesis that the length of the G_4_C_2_ repeat is an important modifier of age of onset and disease risk, we established an allelic series of mice from our most penetrant BAC transgenic line (C9-500). Taking advantage of intergenerational repeat instability observed during the maintenance of our colony (Liu et al., 2016), we selected and bred C9-500 animals that had repeat contractions or expansions and established sublines with 50 (C9-50) or 800 (C9-800) repeats (Fig 4A, B). Limited somatic repeat instability was observed between tail and brain DNA from the 50 and 800 lines (Fig 4B). The limited somatic instability of the G_4_C_2_•G_2_C_4_ expansion, the single-copy of the *C9orf72* transgene and the identical insertion site shared by the C9-800, C9-500, and C9-50 sublines make these mice an ideal resource to test the effects of repeat length on age of onset and disease penetrance.

**Figure 4:**
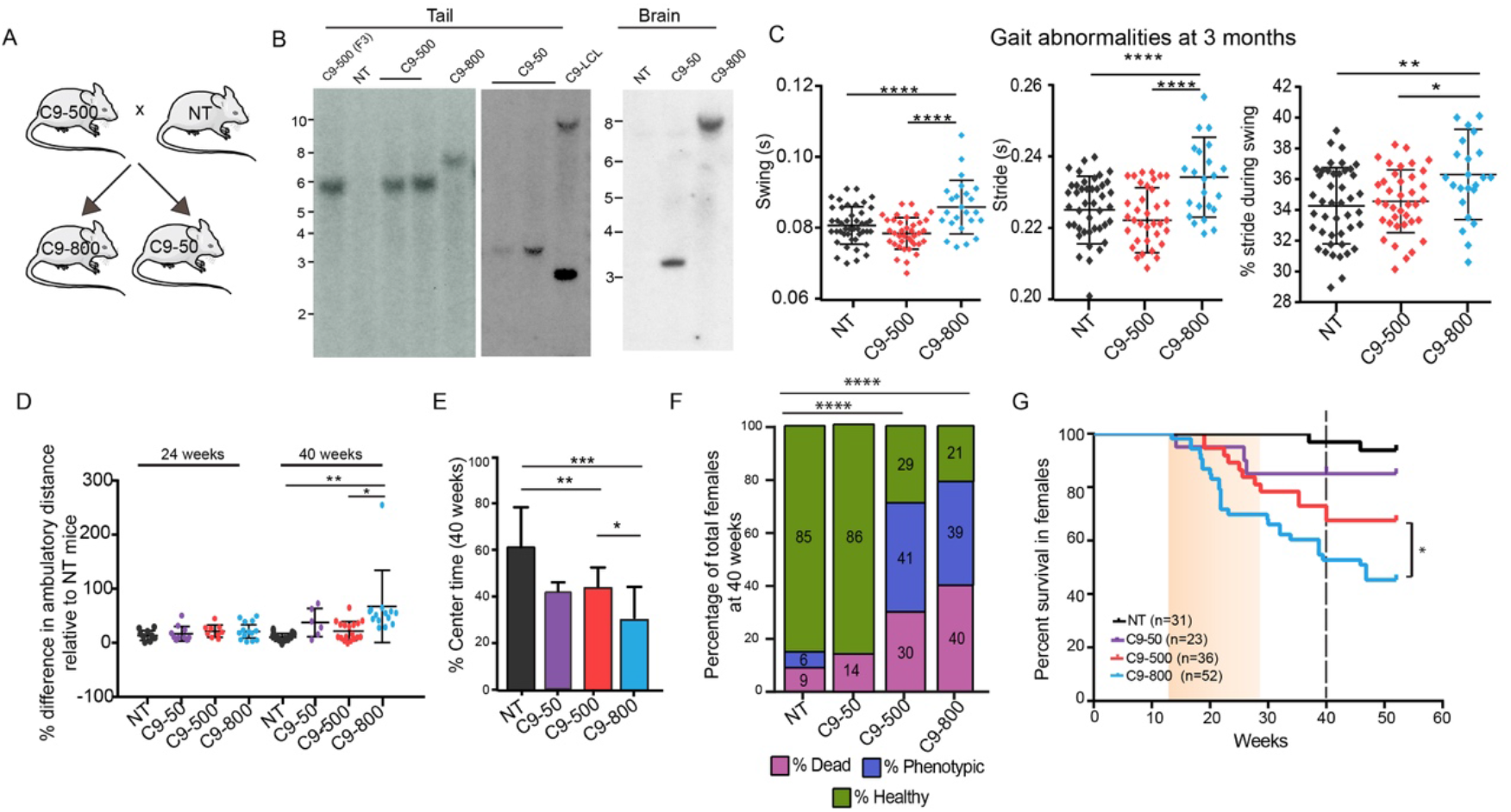
Earlier onset and increased penetrance in isogenic C9 sublines with longer G4C2 expansions. A) Breeding strategy of expansion and contraction lines established from the C9-500 animals with spontaneous intergenerational instability. B) Southern blots of tail and brain DNA from F3 C9-500 mice and C9-800 and C9-50 sublines. C) Gait abnormalities in C9-800 mice at 3 months of age. NT n=44, C9-500 n=38, C9-800 n=24. All parameters listed in Table 3. D) Open field analyses comparing the percentage change in ambulatory distance in C9-50, C9-500 and C9-800 mice compared to NT mice at 24 weeks (NT n=9, C9-50=10, C9-500=9, C9-800=13) and 40 weeks (NT n=9, C9-50=6, C9-500=18, C9-800=10). E) Open field analyses showing percentage time spent in center of chamber for NT, C9-50, C9-500, C9-800 cohorts (NT n=9, C9-50=6, C9-500=18, C9-800=10). F) Population census with percentage of female mice that are sick, healthy and phenotypic using multifactorial scoring criteria. G) Kaplan-Meier survival curves of female mice from NT (n=31), C9-50 (n=23), C9-500 (n=36) and C9-800 (n=52) cohorts. Data information: Statistical analyses for panels D and E were performed using one-way ANOVA and Tukey’s multiple comparison test with mean +/− SEM shown; not significant (ns) = p > 0.05; **p<0.01; ****p <0.0001; ****p <0.0001. In (F), Chi square test was used, ****p<0.0001. In (G) Gehan-Breslow-Wilcoxon test, *p<0.05.

To understand the effects of repeat length on disease, we performed a series of experiments on female mice from this allelic series. First, we performed DigiGait analyses at an early timepoint to test if phenotypes in the C9-800 mice were detected earlier than in the C9-500 cohort. At 12 weeks of age, C9-800 mice showed abnormalities in 9 of 42 DigiGait parameters while the C9-500 mice showed 6 differences compared to NT mice (Table 3). Additionally, 12 parameters were different between C9-800 and C9-500 mice (Fig 4C) including three key parameters typically involved in ALS: swing time, stride time and time of a stride when paw is in swing motion (Mancuso, Olivan et al., 2011, Wooley, Sher et al., 2005). These data show gait abnormalities are found earlier in mice with longer repeats.

**Table 3:** DigiGait parameters of 12-week old females.

| Parameters (41 in total) | Unit | NT vs C9-500 | NT vs C9-800 | C9-500 vs C9-800 |
| --- | --- | --- | --- | --- |
| **Significant Parameters**<br>**(Desired FDR=5%)** | **(real#)** | **6** | **9** | **12** |
| Swing | (s) | ns | ****p<0.0001 | ****p<0.0001 |
| Swing/Stride | (%) | ns | **p=0.0052 | *p=0.0176 |
| Brake | (s) | ns | ns | ns |
| Brake/Stride | (%) | ns | ns | ns |
| Propel | (s) | ns | ns | *p=0.0255 |
| Propel/Stride | (%) | ns | ns | ns |
| Stance | (s) | *p=0.0444 | ns | **p=0.0020 |
| Stance/Stride | (%) | ns | **p=0.0052 | *p=0.0174 |
| Stride | (s) | ns | ****p<0.0001 | ****p<0.0001 |
| Brake/Stance | (%) | ns | ns | ns |
| Propel/Stance | (%) | ns | ns | ns |
| Stance/Swing | (real#) | ns | *p=0.0134 | ns |
| Stride Length | (cm) | ns | ****p<0.0001 | ****p<0.0001 |
| Stride Frequency | (steps/s) | *p=0.0292 | ***p=0.0001 | ****p<0.0001 |
| Paw Angle | (deg) | ns | ns | ns |
| Absolute Paw Angle | (deg) | ns | ns | ns |
| Paw Angle Variability | (deg) | ns | ns | ns |
| Stance Width | (cm) | *p=0.0292 | ns | ns |
| Step Angle | (deg) | ns | ns | ns |
| Stride Length Variability | (cm) | ns | ns | ns |
| Stance Width Variability | (cm) | **p=0.059 | ns | ns |
| Step angle Variability | (deg) | ns | ns | *p=0.0283 |
| #Steps | (real#) | ns | ns | ns |
| Stride Length CV | (CV%) | ****p<0.0001 | ****p<0.0001 | ns |
| Stance Width CV | (CV%) | ***p=0.0006 | ns | *p=0.0243 |
| Step Angle CV | (CV%) | ns | ns | *p=0.0202 |
| Swing Duration CV | (CV%) | ns | ns | ns |
| Paw Area at Peak Stance | (cm<sup>2</sup>) | ns | ns | ns |
| Paw Area Variability at Peak Stance | (cm<sup>2</sup>) | ns | ns | ns |
| Hind Limb Shared Stance Time | (s) | ns | ns | ns |
| Shared/Stance | (%) | ns | ns | ns |
| Stance Factor | (real#) | ns | ns | ns |
| Gait Symmetry | (real#) | ns | ***p=0.0001 | ****p<0.0001 |
| MAX dA/dT | (cm<sup>2</sup>/s) | ns | ns | ns |
| MIN dA/dT | (cm<sup>2</sup>/s) | ns | ns | ns |
| Tau - Propulsion | (real#) | ns | ns | ns |
| Overlap Distance | (cm) | ns | ns | ns |
| Paw Placement Positioning | (cm) | ns | ns | ns |
| Ataxia Coefficient | (real#) | ns | ns | ns |
| Midline Distance | (cm) | ns | ns | ns |
| Paw Drag | (mm<sup>2</sup>) | ns | ns | ns |

#, number; CV, coefficient of variation; ns, not significant in one way ANOVA; desired FDR (false discovery rate) set to 5%; NT n=44; C9-500 n=38; C9-800 n=24.

**Table 4:**
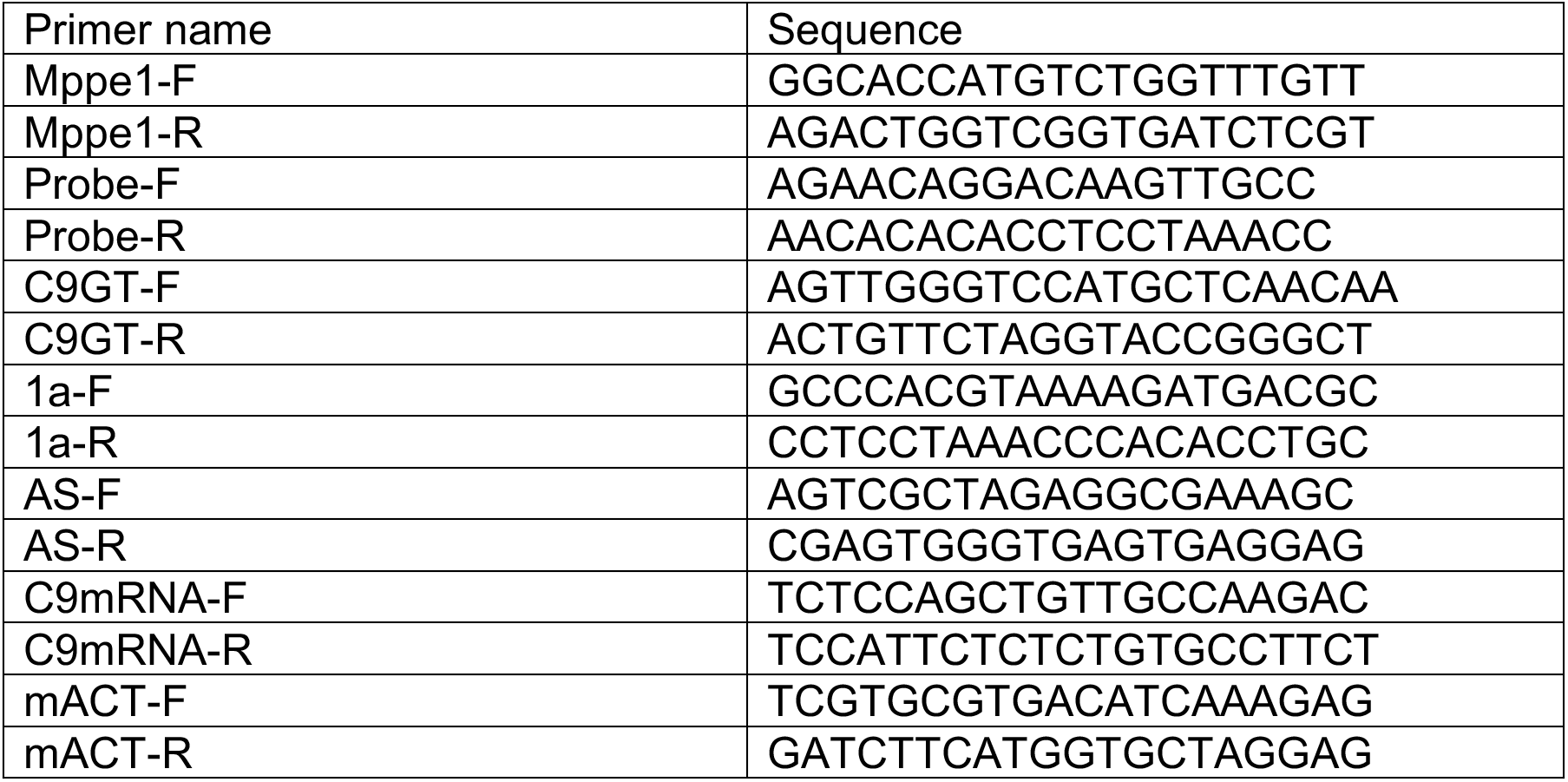
Primers used for PCR.

Open field studies previously showed both slowed movement and hyperactivity in old C9-500 mice (12-18 months) (Liu et al., 2016). To test if increased repeat length causes earlier open field abnormalities, we examined the allelic series of mice at 24 and 40 weeks of age. At 24 weeks there were no differences in ambulatory distance between any of the C9-50, C9-500 and C9-800 sublines compared to NT animals. In contrast, at 40 weeks the C9-800 group showed a significant increase in ambulatory distance compared to both the C9-500 and NT cohorts (Fig 4D). In contrast, no significant differences were found between the C9-500 or C9-50 animals and NT controls at this age. Open field analyses were also used to measure decreases in center time, a phenotype associate with anxiety-like behavior. At 40 weeks, both the C9-500 and C9-800 animals showed decreased center time compared to NT controls. Additionally, center time in the C9-800 animals was decreased compared to C9-500 mice (Fig 4E). Taken together, these data show open field abnormalities are found earlier with increased repeat length.

Phenotypic mice show abnormal cage behavior including kyphosis, inactivity, severe dehydration, and hind-limb paralysis. Population census and cage behavior analyses, performed as previously described (Liu et al., 2016), comparing the percentages of dead, phenotypic and apparently healthy animals at 40 weeks, show significant differences between the C9-500 and C9-800 cohorts compared to NT controls. In contrast, C9-50 animals were not significantly different from NT animals (Fig 4F). Although the overall phenotype distributions in the C9-50 cohort were not significantly different from the NT mice, several mice in the C9-50 line died with features of ALS/FTD including paralysis, kyphosis, weight loss and neurodegeneration. These data demonstrate that repeats as short as 50 can cause C9orf72 ALS/FTD phenotypes, but with reduced penetrance compared to the C9-500 and C9-800 lines. As previously reported (Liu et al., 2016), death in the C9-500 line begins at ~20 weeks of age. In contrast, death of animals in the C9-800 line begins earlier, at approximately 12 weeks of age. Kaplan Meier analyses shows a significant decrease in survival in the C9-800 line compared to C9-500 animals by 52 weeks (Fig 4G). Taken together, these data demonstrate that increased repeat length leads to increased penetrance and earlier disease onset.

### RNA foci and RAN protein aggregates increase with increased repeat length in C9-BAC mice

Next, we performed experiments to understand the molecular changes associated with increases in repeat length. At 20 weeks of age, there was a significant increase in sense and antisense foci in the C9-800 dentate gyrus compared to the C9-500 line (Fig 5A, B). Similar to our previously published data on mice with shorter repeat tracts (29-37 repeats) (Liu et al., 2016), sense and antisense RNA foci were not detected in animals in the C9-50 line (Fig 5A, B). Next, we compared RAN protein aggregates in the retrosplenial cortex by immunohistochemistry (IHC) or immunofluorescence (IF) using human α-GA_1_ and α-GP_1_ antibodies (Nguyen, Montrasio et al., 2020) (Fig 5C, S8A). At 40 weeks of age, there was a significant increase in percentage of cells with polyGA aggregates (71% vs 40%) and polyGP aggregates (22% vs. 15%) detected by IHC or IF in the C9-800 compared to the C9-500 mice (Fig 5C-E) while no GA or GP aggregates were detected in the C9-50 sub-line consistent with the decreased penetrance in this sub-line. IF studies performed at 20 weeks showed similar trends with significantly higher levels of aggregates between the C9-800 vs. NT and C9-500 vs. NT but no difference between these groups (Fig S8B, C). The levels of soluble GP protein measured by MSD showed similar trends at 40 weeks (Fig S8D).

**Figure 5:**
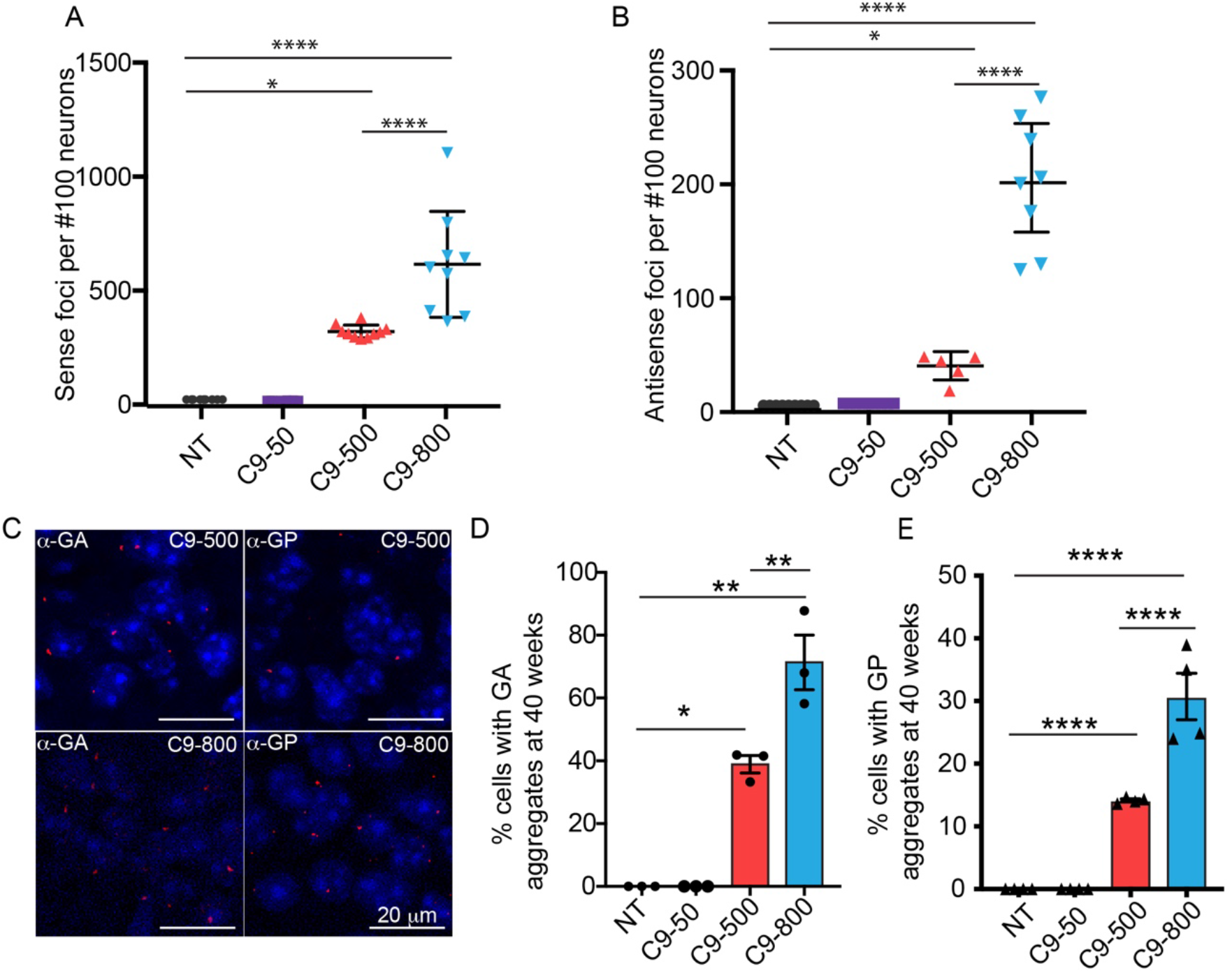
Molecular features of C9orf72 ALS/FTD in C9-BAC mice increase with increase in repeat length. A, B) Quantification of fluorescence in situ hybridization (FISH) detection of sense (A) and antisense (B) RNA foci in the dentate gyrus of the hippocampus. C) Immunofluorescence (IF) showing representative images of GA and GP RAN protein aggregates in the retrosplenial cortex of 40-week old female C9-500 and C9-800 mice. D) Quantification of GA and GP RAN protein aggregates was done in a blinded fashion and is shown as the percentage of neurons with aggregates in NT, C9-50, C9-500 and C9-800 mouse cohorts. Data information: Statistical analyses for (A), (B) and (D) done using a one-way ANOVA with Bonferroni correction for multiple comparisons. Mean +/− SEM, *p<0.05, **p<0.01, ****p <0.0001. See also Figure S8.

In summary, our data demonstrate that repeat length is a modifier of disease in our C9-BAC transgenic mouse model. Mice with longer G_4_C_2_ repeat tracts show increased disease penetrance and earlier ages of onset and are characterized by increased levels of RNA foci and RAN protein aggregates.

## Discussion

To understand the complex molecular mechanisms of *C9orf72* ALS/FTD, more than a dozen mouse models have been developed (Atanasio et al., 2016, Burberry et al., 2016, Chew et al., 2015, Hao et al., 2019, Jiang & Cleveland, 2016, Liu et al., 2016, O’Rourke et al., 2015, Peters et al., 2015, Schludi et al., 2017, Zhang et al., 2018, Zhang et al., 2019). Among these, we reported multiple lines of *C9orf72* ALS/FTD BAC transgenic mice that develop the behavioral, neuropathological and molecular features of disease (Liu et al., 2016). To further understand disease in this mouse model, we performed whole genome sequencing and show four independent lines of these BAC mice have unique integration sites. RNA sequencing identified alternative splicing changes affecting RNA processing and degradation pathways as an early molecular signature of disease, which worsens with disease progression. Similar to *C9orf72* ALS/FTD patients (Butti & Patten, 2018, Prudencio et al., 2015, Prudencio et al., 2017), gene expression changes in neuroinflammatory and neurodegenerative pathways predominate in severely affected end-stage animals. Finally, isogenic sub-lines generated from the single copy C9-500 line with 800 repeats (C9-800) show increased RNA foci and RAN protein aggregates as well as earlier ages of onset and increased disease penetrance. These data demonstrate that phenotypes in our C9-BAC mouse model are caused by the repeat expansion, occur independent of transgene integration site and that longer repeat lengths decrease age of onset and increase disease penetrance.

Patients with C9orf72 expansions mutation show remarkable heterogeneity in clinical presentation (Van Mossevelde et al., 2017a, Van Mossevelde et al., 2017b) and age of onset (Chio et al., 2012, DeJesus-Hernandez et al., 2011, Gijselinck et al., 2012, Gusella & MacDonald, 2009, Paulson, 2018, Renton et al., 2011, Stewart et al., 2012, van der Zee et al., 2013, Van Langenhove et al., 2013, Van Mossevelde et al., 2017a), but the role of repeat length as a disease modifier has been difficult to determine for a number of reasons. First, there is substantial somatic repeat instability and repeat lengths in blood are likely to be substantially shorter than those found in affected brain tissue (Nordin, Akimoto et al., 2015, van Blitterswijk, Baker et al., 2013). Southern blotting is the most common way to measure repeat length but somatic instability and the variation of repeat lengths within a single individual make comparisons of repeat length and age of onset in patients inaccurate. Additionally, GC rich expansion mutations are difficult to amplify by PCR and to sequence making small-pool PCR strategies to accurately measure the distributions of repeat lengths challenging (Dandelot & Gourdon, 2018, Gomes-Pereira, Bidichandani et al., 2004).

Our generation of an isogenic series of C9-BAC mice with 800, 500 or 50 repeats has allowed us to directly demonstrate that longer repeat tracts result in earlier onset of behavioral abnormalities and decreased survival compared to shorter alleles. These results, combined with the somatic instability seen in patients, suggest that targeting DNA repair pathways may be a viable approach for mitigating disease in C9orf72 ALS/FTD (Jones et al., 2017, Lee et al., 2015, Wright et al., 2019). While reducing repeat tract length in our mice did reduce disease penetrance, several C9-50 mice did show ALS/FTD phenotypes. The phenotypes found in the C9-50 animals, combined with previously published phenotypes in animals with 4 copies of the transgene containing 36-29 repeats (Liu et al., 2016), suggest additional human studies are needed to understand the relative risks of shorter repeat expansion mutations and the potential role of somatic instability to exacerbate disease. This may be particularly relevant for families with sporadic cases of *C9orf72* ALS/FTD.

Ectopic cytoplasmic localization of mutant TDP43, FUS and MATR3 or overexpression of wildtype TDP43, FUS and MATR3 protein has been shown to be associated with ALS phenotypes in patients and model systems, respectively (Lagier-Tourenne, Polymenidou et al., 2010, Malik, Miguez et al., 2018, Mitchell, McGoldrick et al., 2013, Tada, Doi et al., 2018, Wils, Kleinberger et al., 2010). C9orf72, a DENN (differentially expressed in normal and neoplastic cells) domain containing protein, has been shown to play a role in autophagy and immune-regulatory functions (Lall & Baloh, 2017, Levine, Daniels et al., 2013, Nassif, Woehlbier et al., 2017). Recently, it has been shown that a dose dependent increase in motor deficits was observed when single copy C9-500 mice were crossed with C9orf72 heterozygous and homozygous mice suggesting both loss of function and gain of function effects may contribute to C9orf72 ALS/FTD (Shao, Liang et al., 2019). Additionally, it has been proposed that overexpression of C9orf72 protein may lead to toxicity in mouse models of disease (Hayes & Rothstein, 2016). Our data, showing that there are no significant differences in C9orf72 protein levels between C9-500 and NT mice combined with our allelic series data, indicate that repeat length and not C9orf72 overexpression is the primary driver of the ALS/FTD phenotypes found in our mice.

Substantial data suggest RNA gain-of-function effects contribute to *C9orf72* ALS/FTD (Barmada, 2015; Scotti and Swanson, 2016; Taylor et al., 2016) and a large number of RNA binding proteins have been proposed to be sequestered by the repeats. (Haeusler et al., 2016; Taylor et al., 2016). Circular dichroism studies performed on short stretches of sense repeats G4C2 show that the repeat can adopt G quadruplex, R loop or hairpin conformations and the antisense repeats likely adopt a hairpin conformation (Haeusler et al., 2014, Mizielinska, Grönke et al., 2014, Reddy, Zamiri et al., 2013). Since several of these predicted RNA binding proteins also contain low complexity domains, it is possible that multiple RNA binding proteins interact with repeat RNAs and form dynamic liquid droplet like structures that may be complicated to resolve (Jain & Vale, 2017, Lee, Zhang et al., 2016, Lin, Mori et al., 2016). The variability and complexity in secondary structures formed by the repeats make it more likely that multiple RNA binding protein are sequestered by these repeat motifs (Cooper-Knock et al., 2014), a possibility consistent with our data showing that genes with abnormal splicing events have a variety of RNA recognition motifs.

Transcriptomic data from our acute C9-BAC mice are consistent with gene expression changes caused by ongoing apoptotic processes involved in cell death and inflammation found at end-stage disease. In contrast, transcriptomic profiles of presymptomatic mice showed minimal changes in gene expression but abundant changes in alternative splicing. Robust alternative splicing changes are also detected in neurons differentiated from C9orf72 patient iPSCs (iPSN), and similarly showed few gene expression differences compared to C9orf72 patient autopsy tissue (Donnelly et al., 2013). In our mice, 240 genes are misspliced in C9(+) presymptomatic animals, and the psi values of 83 of these genes increase further in mice with acute end-stage disease. These genes include *Elavl2*, which has an RNA recognition motif (RRM) and has been predicted to play a role in ALS because of its similarity to TDP-43 and FUS (Lu, Lim et al., 2017). Pard3, which regulates neuronal polarity has also been shown to be mis-spliced in ALS patient autopsy tissue (Yin, Lopez-Gonzalez et al., 2017b). These results, combined with the 83 other alternative spliced genes whose psi values increase with disease progression, highlight these alternative splicing changes as a novel and early molecular signature of *C9orf72* ALS/FTD.

A role of RAN proteins in disrupting alternative splicing has also been proposed. In cell culture experiments, ~5000 mis-spliced events were observed when astrocytes were treated with PR RAN protein (Kwon, Xiang et al., 2014). Additionally, it has been shown that in cell culture systems, GR and PR RAN proteins associate with low complexity domains of RNA binding proteins (Lee et al., 2016, Lin et al., 2016, Shi, Mori et al., 2017) and can cause mis-splicing in a U2snRNP dependent manner (Yin, Lopez-Gonzalez et al., 2017a). In cell culture, these low complexity domain proteins are typically involved in the formation of membrane-less organelles such as stress granules and neuronal speckles (Protter & Parker, 2016, Taylor, Brown et al., 2016). Since splicing factors normally localize to the neuronal speckles, it is possible that RAN proteins interact with RNA binding proteins containing low complexity domains and disrupt splicing globally. Global splicing deficits would be likely to cause an increase in overall alternative spliced changes rather than the disruption of specific set of events mediated by the sequestration of a single family of RNA binding protein as is seen in myotonic dystrophy (Scotti & Swanson, 2016). These data combined with our data showing phenotypic C9-500 mice have more RAN protein aggregates compared to asymptomatic C9-500 (Liu et al., 2016), suggests that RAN proteins could contribute to the early RNA splicing abnormalities found in our mice.

Longer repeat lengths result in the expression of both longer expansion RNAs and increased RAN protein aggregates. Since both RNA GOF and protein toxicity are closely connected to repeat length, additional work will be needed to clarify the relative roles of RNA gain of function effects and RAN proteins in disease. Our mouse model is an excellent platform to address the contribution of specific disease mechanisms and for the development of potential therapies. These C9-BAC mice were recently used to demonstrate that a human antibody targeting the GA RAN protein is sufficient to rescue behavior and survival phenotypes, without changing the levels of their corresponding expansion RNAs. These preclinical data highlight the therapeutic potential of targeting RAN proteins (Nguyen et al., 2020).

Since the discovery of the *C9orf72* expansion mutation, research has focused on understanding disease mechanisms and identifying therapeutic targets. Transgenic mouse models have contributed to these efforts, but ectopic overexpression and transgene integration can affect the resulting phenotypes and suitability for testing potential therapies. This issue is not restricted to C9-ALS as this issue also occurs in the widely used R6/2 mouse model of Huntington disease (Jacobsen, Erdin et al., 2017). Multiple features make our C9-500 BAC transgenic mice a valuable tool both for understanding the biology and time course of the disease and also for testing novel therapeutic strategies. First, the phenotypes in our mice, which contain the full-length *C9orf72* gene and substantial flanking sequences, are not affected by integration site. Second, the C9-500 line carries one copy of the transgene expressed at levels comparable to the endogenous *C9orf72* ortholog in mice. Third, we have identified a group of alternative splicing as a molecular signature of disease that worsen with disease progression and may be useful biomarkers of disease. Finally, we demonstrate that repeat length increases disease penetrance and that the phenotypes in our mice are caused by a gain of function of the repeat expansion. Taken together, our data provide mechanistic insight into *C9orf72* ALS/FTD and demonstrate that our mice are a robust tool to study the disease and to test therapeutic strategies for disease intervention.

## Acknowledgments

**Non-author contributions:** We thank Dr. John Cleary, Dr. Yuanjing Liu and Dr. Maurice Swanson for input and advice on the manuscript, Dr. Fabio Montrasio and Dr. Jan Grimm for providing human antibodies for polyGA and polyGP detection.

**Funding:** We thank the National Institutes of Health (R01 NS098819), Target ALS the ALS Association, the Packard Center and the Muscular Dystrophy Association for support

## Author Contributions

*<u>Performed experiments</u>*: A. Pattamatta, L. Nguyen, H. Olafson, M. Scotti, L. A. Laboissonniere, J. Richardson. *<u>Provided reagents:</u>* J. Berglund, E. Wang, L. Ranum, Analyzed data: A. Pattamatta, L. Nguyen, H. Olafson, T. Zu, E.Wang, L. Ranum. *<u>Data and material availability:</u>* Raw data and reagents will be available upon request from Dr. Ranum. *<u>Wrote and revised the manuscript:</u>* A. Pattamatta and L. Ranum wrote the manuscript, A. Pattamatta, L. Nguyen and L. Ranum revised the manuscript with input from all the authors.

## Conflict of Interest

The authors declare that they have no conflict of interest.

## Notes

### Competing Interest Statement

The authors have declared no competing interest.

## References

Ash PE, Bieniek KF, Gendron TF, Caulfield T, Lin WL, Dejesus-Hernandez M, van Blitterswijk MM, Jansen-West K, Paul JW, 3rd, Rademakers R, Boylan KB, Dickson DW, Petrucelli L (2013) Unconventional translation of C9ORF72 GGGGCC expansion generates insoluble polypeptides specific to c9FTD/ALS. Neuron 77: 639–46

Atanasio A, Decman V, White D, Ramos M, Ikiz B, Lee HC, Siao CJ, Brydges S, LaRosa E, Bai Y, Fury W, Burfeind P, Zamfirova R, Warshaw G, Orengo J, Oyejide A, Fralish M, Auerbach W, Poueymirou W, Freudenberg J et al. (2016) C9orf72 ablation causes immune dysregulation characterized by leukocyte expansion, autoantibody production, and glomerulonephropathy in mice. Sci Rep 6: 23204

Bray NL, Pimentel H, Melsted P, Pachter L (2016) Near-optimal probabilistic RNA-seq quantification. Nat Biotechnol 34: 525–7

Burberry A, Suzuki N, Wang JY, Moccia R, Mordes DA, Stewart MH, Suzuki-Uematsu S, Ghosh S, Singh A, Merkle FT, Koszka K, Li QZ, Zon L, Rossi DJ, Trowbridge JJ, Notarangelo LD, Eggan K (2016) Loss-of-function mutations in the C9ORF72 mouse ortholog cause fatal autoimmune disease. Sci Transl Med 8: 347ra93

Butti Z, Patten SA (2018) RNA Dysregulation in Amyotrophic Lateral Sclerosis. Front Genet 9: 712

Chew J, Gendron TF, Prudencio M, Sasaguri H, Zhang YJ, Castanedes-Casey M, Lee CW, Jansen-West K, Kurti A, Murray ME, Bieniek KF, Bauer PO, Whitelaw EC, Rousseau L, Stankowski JN, Stetler C, Daughrity LM, Perkerson EA, Desaro P, Johnston A et al. (2015) Neurodegeneration. C9ORF72 repeat expansions in mice cause TDP-43 pathology, neuronal loss, and behavioral deficits. Science 348: 1151–4

Chio A, Restagno G, Brunetti M, Ossola I, Calvo A, Canosa A, Moglia C, Floris G, Tacconi P, Marrosu F, Marrosu MG, Murru MR, Majounie E, Renton AE, Abramzon Y, Pugliatti M, Sotgiu MA, Traynor BJ, Borghero G, Consortium S (2012) ALS/FTD phenotype in two Sardinian families carrying both C9ORF72 and TARDBP mutations. J Neurol Neurosurg Psychiatry 83: 730–3

Conlon EG, Lu L, Sharma A, Yamazaki T, Tang T, Shneider NA, Manley JL (2016) The C9ORF72 GGGGCC expansion forms RNA G-quadruplex inclusions and sequesters hnRNP H to disrupt splicing in ALS brains. Elife 5

Cooper-Knock J, Walsh MJ, Higginbottom A, Robin Highley J, Dickman MJ, Edbauer D, Ince PG, Wharton SB, Wilson SA, Kirby J, Hautbergue GM, Shaw PJ (2014) Sequestration of multiple RNA recognition motif-containing proteins by C9orf72 repeat expansions. Brain 137: 2040–51

Dandelot E, Gourdon G (2018) The flash-small-pool PCR: how to transform blotting and numerous hybridization steps into a simple denatured PCR. Biotechniques 64: 262–265

DeJesus-Hernandez M, Mackenzie IR, Boeve BF, Boxer AL, Baker M, Rutherford NJ, Nicholson AM, Finch NA, Flynn H, Adamson J, Kouri N, Wojtas A, Sengdy P, Hsiung GY, Karydas A, Seeley WW, Josephs KA, Coppola G, Geschwind DH, Wszolek ZK et al. (2011) Expanded GGGGCC hexanucleotide repeat in noncoding region of C9ORF72 causes chromosome 9p-linked FTD and ALS. Neuron 72: 245–56

Dobin A, Davis CA, Schlesinger F, Drenkow J, Zaleski C, Jha S, Batut P, Chaisson M, Gingeras TR (2013) STAR: ultrafast universal RNA-seq aligner. Bioinformatics 29: 15–21

Donnelly CJ, Zhang PW, Pham JT, Heusler AR, Mistry NA, Vidensky S, Daley EL, Poth EM, Hoover B, Fines DM, Maragakis N, Tienari PJ, Petrucelli L, Traynor BJ, Wang JO, Rigo F, Bennett CF, Blackshaw S, Sattler R, Rothstein JD (2013) RNA Toxicity from the ALS/FTD C9ORF72 Expansion Is Mitigated by Antisense Intervention. Neuron 80: 415–428

Ebbert MTW, Ross CA, Pregent LJ, Lank RJ, Zhang C, Katzman RB, Jansen-West K, Song YP, da Rocha EL, Palmucci C, Desaro P, Robertson AE, Caputo AM, Dickson DW, Boylan KB, Rademakers R, Ordog T, Li H, Belzil VV (2017) Conserved DNA methylation combined with differential frontal cortex and cerebellar expression distinguishes C9orf72-associated and sporadic ALS, and implicates SERPINA1 in disease. Acta Neuropathologica 134: 715–728

Freibaum BD, Lu Y, Lopez-Gonzalez R, Kim NC, Almeida S, Lee KH, Badders N, Valentine M, Miller BL, Wong PC, Petrucelli L, Kim HJ, Gao FB, Taylor JP (2015) GGGGCC repeat expansion in C9orf72 compromises nucleocytoplasmic transport. Nature 525: 129–33

Gijselinck I, Van Langenhove T, van der Zee J, Sleegers K, Philtjens S, Kleinberger G, Janssens J, Bettens K, Van Cauwenberghe C, Pereson S, Engelborghs S, Sieben A, De Jonghe P, Vandenberghe R, Santens P, De Bleecker J, Maes G, Baumer V, Dillen L, Joris G et al. (2012) A C9orf72 promoter repeat expansion in a Flanders-Belgian cohort with disorders of the frontotemporal lobar degeneration-amyotrophic lateral sclerosis spectrum: a gene identification study. Lancet Neurol 11: 54–65

Gomes-Pereira M, Bidichandani SI, Monckton DG (2004) Analysis of unstable triplet repeats using small-pool polymerase chain reaction. Methods Mol Biol 277: 61–76

Gusella JF, MacDonald ME (2000) Molecular genetics: unmasking polyglutamine triggers in neurodegenerative disease. Nat Rev Neurosci 1: 109–15

Gusella JF, MacDonald ME (2009) Huntington’s disease: the case for genetic modifiers. Genome Medicine 1

Haeusler AR, Donnelly CJ, Periz G, Simko EA, Shaw PG, Kim MS, Maragakis NJ, Troncoso JC, Pandey A, Sattler R, Rothstein JD, Wang J (2014) C9orf72 nucleotide repeat structures initiate molecular cascades of disease. Nature 507: 195–200

Hao ZB, Liu L, Tao ZT, Wang R, Ren HG, Sun HY, Lin ZX, Zhang ZX, Mu CC, Zhou JW, Wang GH (2019) Motor dysfunction and neurodegeneration in a C9orf72 mouse line expressing poly-PR. Nat Commun 10

Hayes LR, Rothstein JD (2016) C9ORF72-ALS/FTD: Transgenic Mice Make a Come-BAC. Neuron 90: 427–31

Hsiung GY, DeJesus-Hernandez M, Feldman HH, Sengdy P, Bouchard-Kerr P, Dwosh E, Butler R, Leung B, Fok A, Rutherford NJ, Baker M, Rademakers R, Mackenzie IR (2012) Clinical and pathological features of familial frontotemporal dementia caused by C9ORF72 mutation on chromosome 9p. Brain 135: 709–22

Ince-Dunn G, Okano HJ, Jensen KB, Park WY, Zhong R, Ule J, Mele A, Fak JJ, Yang C, Zhang C, Yoo J, Herre M, Okano H, Noebels JL, Darnell RB (2012) Neuronal Elav-like (Hu) proteins regulate RNA splicing and abundance to control glutamate levels and neuronal excitability. Neuron 75: 1067–80

Jacobsen JC, Erdin S, Chiang C, Hanscom C, Handley RR, Barker DD, Stortchevoi A, Blumenthal I, Reid SJ, Snell RG, MacDonald ME, Morton AJ, Ernst C, Gusella JF, Talkowski ME (2017) Potential molecular consequences of transgene integration: The R6/2 mouse example. Sci Rep 7: 41120

Jain A, Vale RD (2017) RNA phase transitions in repeat expansion disorders. Nature 546: 243–247

Jiang J, Cleveland DW (2016) Bidirectional Transcriptional Inhibition as Therapy for ALS/FTD Caused by Repeat Expansion in C9orf72. Neuron 92: 1160–1163

Jiang J, Zhu Q, Gendron TF, Saberi S, McAlonis-Downes M, Seelman A, Stauffer JE, Jafar-Nejad P, Drenner K, Schulte D, Chun S, Sun S, Ling SC, Myers B, Engelhardt J, Katz M, Baughn M, Platoshyn O, Marsala M, Watt A et al. (2016) Gain of Toxicity from ALS/FTD-Linked Repeat Expansions in C9ORF72 Is Alleviated by Antisense Oligonucleotides Targeting GGGGCC-Containing RNAs. Neuron 90: 535–50

Jones L, Houlden H, Tabrizi SJ (2017) DNA repair in the trinucleotide repeat disorders. Lancet Neurology 16: 88–96

Jovicic A, Mertens J, Boeynaems S, Bogaert E, Chai N, Yamada SB, Paul JW, 3rd, Sun S, Herdy JR, Bieri G, Kramer NJ, Gage FH, Van Den Bosch L, Robberecht W, Gitler AD (2015) Modifiers of C9orf72 dipeptide repeat toxicity connect nucleocytoplasmic transport defects to FTD/ALS. Nat Neurosci 18: 1226–9

Kwon I, Xiang S, Kato M, Wu L, Theodoropoulos P, Wang T, Kim J, Yun J, Xie Y, McKnight SL (2014) Poly-dipeptides encoded by the C9orf72 repeats bind nucleoli, impede RNA biogenesis, and kill cells. Science 345: 1139–45

Laflamme C, McKeever PM, Kumar R, Schwartz J, Kolahdouzan M, Chen CX, You Z, Benaliouad F, Gileadi O, McBride HM, Durcan TM, Edwards AM, Healy LM, Robertson J, McPherson PS (2019) Implementation of an antibody characterization procedure and application to the major ALS/FTD disease gene C9ORF72. Elife 8

Lagier-Tourenne C, Polymenidou M, Cleveland DW (2010) TDP-43 and FUS/TLS: emerging roles in RNA processing and neurodegeneration. Hum Mol Genet 19: R46–64

Lall D, Baloh RH (2017) Microglia and C9orf72 in neuroinflammation and ALS and frontotemporal dementia. J Clin Invest 127: 3250–3258

Lee JM, Wheeler VC, Chao MJ, Vonsattel JPG, Pinto RM, Lucente D, Abu-Elneel K, Ramos EM, Mysore JS, Gillis T, MacDonald ME, Gusella JF, Harold D, Stone TC, Escott-Price V, Han J, Vedernikov A, Holmans P, Jones L, Kwak S et al. (2015) Identification of Genetic Factors that Modify Clinical Onset of Huntington’s Disease. Cell 162: 516–526

Lee KH, Zhang P, Kim HJ, Mitrea DM, Sarkar M, Freibaum BD, Cika J, Coughlin M, Messing J, Molliex A, Maxwell BA, Kim NC, Temirov J, Moore J, Kolaitis RM, Shaw TI, Bai B, Peng J, Kriwacki RW, Taylor JP (2016) C9orf72 Dipeptide Repeats Impair the Assembly, Dynamics, and Function of Membrane-Less Organelles. Cell 167: 774–788.e17

Lee YB, Chen HJ, Peres JN, Gomez-Deza J, Attig J, Stalekar M, Troakes C, Nishimura AL, Scotter EL, Vance C, Adachi Y, Sardone V, Miller JW, Smith BN, Gallo JM, Ule J, Hirth F, Rogelj B, Houart C, Shaw CE (2013) Hexanucleotide repeats in ALS/FTD form length-dependent RNA foci, sequester RNA binding proteins, and are neurotoxic. Cell Rep 5: 1178–86

Levine TP, Daniels RD, Gatta AT, Wong LH, Hayes MJ (2013) The product of C9orf72, a gene strongly implicated in neurodegeneration, is structurally related to DENN Rab-GEFs. Bioinformatics 29: 499–503

Li H (2013) Aligning sequence reads, clone sequences and assembly contigs with BWA-MEM. arXiv:13033997v1 [q-bioGN]

Lin Y, Mori E, Kato M, Xiang S, Wu L, Kwon I, McKnight SL (2016) Toxic PR Poly-Dipeptides Encoded by the C9orf72 Repeat Expansion Target LC Domain Polymers. Cell 167: 789–802.e12

Liu Y, Pattamatta A, Zu T, Reid T, Bardhi O, Borchelt DR, Yachnis AT, Ranum LP (2016) C9orf72 BAC Mouse Model with Motor Deficits and Neurodegenerative Features of ALS/FTD. Neuron 90: 521–34

Lu YM, Lim LZ, Song JX (2017) RRM domain of ALS/FTD-causing FUS characteristic of irreversible unfolding spontaneously selfassembles into amyloid fibrils. Sci Rep-Uk 7

Majounie E, Renton AE, Mok K, Dopper EG, Waite A, Rollinson S, Chio A, Restagno G, Nicolaou N, Simon-Sanchez J, van Swieten JC, Abramzon Y, Johnson JO, Sendtner M, Pamphlett R, Orrell RW, Mead S, Sidle KC, Houlden H, Rohrer JD et al. (2012) Frequency of the C9orf72 hexanucleotide repeat expansion in patients with amyotrophic lateral sclerosis and frontotemporal dementia: a cross-sectional study. Lancet Neurol 11: 323–30

Malik AM, Miguez RA, Li X, Ho YS, Feldman EL, Barmada SJ (2018) Matrin 3-dependent neurotoxicity is modified by nucleic acid binding and nucleocytoplasmic localization. Elife 7

Mancuso R, Olivan S, Osta R, Navarro X (2011) Evolution of gait abnormalities in SOD1(G93A) transgenic mice. Brain Res 1406: 65–73

Marco-Sola S, Sammeth M, Guigo R, Ribeca P (2012) The GEM mapper: fast, accurate and versatile alignment by filtration. Nat Methods 9: 1185–8

Martin M (2011) Cutadapt removes adapter sequences from high-throughput sequencing reads. EMBnetjournal 17: 10–12

McKenna A HM, Banks E, Sivachenko A, Cibulskis K, Kernytsky A, Garimella K, Altshuler D, Gabriel S, Daly M, DePristo MA (2010) The Genome Analysis Toolkit: a MapReduce framework for analyzing next-generation DNA sequencing data. Genome Res 20: 1297–1303

Mitchell JC, McGoldrick P, Vance C, Hortobagyi T, Sreedharan J, Rogelj B, Tudor EL, Smith BN, Klasen C, Miller CC, Cooper JD, Greensmith L, Shaw CE (2013) Overexpression of human wild-type FUS causes progressive motor neuron degeneration in an age- and dose-dependent fashion. Acta Neuropathol 125: 273–88

Mizielinska S, Grönke S, Niccoli T, Ridler CE, Clayton EL, Devoy A, Moens T, Norona FE, Woollacott IO, Pietrzyk J, Cleverley K, Nicoll AJ, Pickering-Brown S, Dols J, Cabecinha M, Hendrich O, Fratta P, Fisher EM, Partridge L, Isaacs AM (2014) C9orf72 repeat expansions cause neurodegeneration in Drosophila through arginine-rich proteins. Science 345: 1192–4

Mizielinska S, Gronke S, Niccoli T, Ridler CE, Clayton EL, Devoy A, Moens T, Norona FE, Woollacott IOC, Pietrzyk J, Cleverley K, Nicoll AJ, Pickering-Brown S, Dols J, Cabecinha M, Hendrich O, Fratta P, Fisher EMC, Partridge L, Isaacs AM (2014) C9orf72 repeat expansions cause neurodegeneration in Drosophila through arginine-rich proteins. Science 345: 1192–1194

Mori K, Weng SM, Arzberger T, May S, Rentzsch K, Kremmer E, Schmid B, Kretzschmar HA, Cruts M, Van Broeckhoven C, Haass C, Edbauer D (2013) The C9orf72 GGGGCC repeat is translated into aggregating dipeptide-repeat proteins in FTLD/ALS. Science 339: 1335–8

Murphy NA, Arthur KC, Tienari PJ, Houlden H, Chio A, Traynor BJ (2017) Age-related penetrance of the C9orf72 repeat expansion. Sci Rep 7: 2116

Nassif M, Woehlbier U, Manque PA (2017) The Enigmatic Role of C9ORF72 in Autophagy. Front Neurosci 11: 442

Nguyen L, Cleary JD, Ranum LPW (2019) Repeat-Associated Non-ATG Translation: Molecular Mechanisms and Contribution to Neurological Disease. Annu Rev Neurosci 42: 227–247

Nguyen L, Montrasio F, Pattamatta A, Tusi SK, Bardhi O, Meyer KD, Hayes L, Nakamura K, Banez-Coronel M, Coyne A, Guo S, Laboissonniere LA, Gu Y, Narayanan S, Smith B, Nitsch RM, Kankel MW, Rushe M, Rothstein J, Zu T et al. (2020) Antibody Therapy Targeting RAN Proteins Rescues C9 ALS/FTD Phenotypes in C9orf72 Mouse Model. Neuron 105: 645–662 e11

Nordin A, Akimoto C, Wuolikainen A, Alstermark H, Jonsson P, Birve A, Marklund SL, Graffmo KS, Forsberg K, Brannstrom T, Andersen PM (2015) Extensive size variability of the GGGGCC expansion in C9orf72 in both neuronal and non-neuronal tissues in 18 patients with ALS or FTD. Hum Mol Genet 24: 3133–42

O’Rourke JG, Bogdanik L, Muhammad AK, Gendron TF, Kim KJ, Austin A, Cady J, Liu EY, Zarrow J, Grant S, Ho R, Bell S, Carmona S, Simpkinson M, Lall D, Wu K, Daughrity L, Dickson DW, Harms MB, Petrucelli L et al. (2015) C9orf72 BAC Transgenic Mice Display Typical Pathologic Features of ALS/FTD. Neuron 88: 892–901

Paulson H (2018) Repeat expansion diseases. Handb Clin Neurol 147: 105–123

Peters OM, Cabrera GT, Tran H, Gendron TF, McKeon JE, Metterville J, Weiss A, Wightman N, Salameh J, Kim J, Sun H, Boylan KB, Dickson D, Kennedy Z, Lin Z, Zhang YJ, Daughrity L, Jung C, Gao FB, Sapp PC et al. (2015) Human C9ORF72 Hexanucleotide Expansion Reproduces RNA Foci and Dipeptide Repeat Proteins but Not Neurodegeneration in BAC Transgenic Mice. Neuron 88: 902–9

Picard A set of command line tools (in Java) for manipulating high-throughput sequencing (HTS) data and formats such as SAM/BAM/CRAM and VCF. http://picard.sourceforge.net.

Protter DSW, Parker R (2016) Principles and Properties of Stress Granules. Trends Cell Biol 26: 668–679

Prudencio M, Belzil VV, Batra R, Ross CA, Gendron TF, Pregent LJ, Murray ME, Overstreet KK, Piazza-Johnston AE, Desaro P, Bieniek KF, DeTure M, Lee WC, Biendarra SM, Davis MD, Baker MC, Perkerson RB, van Blitterswijk M, Stetler CT, Rademakers R et al. (2015) Distinct brain transcriptome profiles in C9orf72-associated and sporadic ALS. Nat Neurosci 18: 1175–82

Prudencio M, Gonzales PK, Cook CN, Gendron TF, Daughrity LM, Song Y, Ebbert MTW, van Blitterswijk M, Zhang YJ, Jansen-West K, Baker MC, DeTure M, Rademakers R, Boylan KB, Dickson DW, Petrucelli L, Link CD (2017) Repetitive element transcripts are elevated in the brain of C9orf72 ALS/FTLD patients. Hum Mol Genet 26: 3421–3431

Reddy K, Zamiri B, Stanley SY, Macgregor RB, Jr., Pearson CE (2013) The disease-associated r(GGGGCC)n repeat from the C9orf72 gene forms tract length-dependent uni- and multimolecular RNA G-quadruplex structures. J Biol Chem 288: 9860–6

Renton AE, Majounie E, Waite A, Simon-Sanchez J, Rollinson S, Gibbs JR, Schymick JC, Laaksovirta H, van Swieten JC, Myllykangas L, Kalimo H, Paetau A, Abramzon Y, Remes AM, Kaganovich A, Scholz SW, Duckworth J, Ding J, Harmer DW, Hernandez DG et al. (2011) A hexanucleotide repeat expansion in C9ORF72 is the cause of chromosome 9p21-linked ALS-FTD. Neuron 72: 257–68

Sareen D, O’Rourke JG, Meera P, Muhammad AK, Grant S, Simpkinson M, Bell S, Carmona S, Ornelas L, Sahabian A, Gendron T, Petrucelli L, Baughn M, Ravits J, Harms MB, Rigo F, Bennett CF, Otis TS, Svendsen CN, Baloh RH (2013) Targeting RNA Foci in iPSC-Derived Motor Neurons from ALS Patients with a C9ORF72 Repeat Expansion. Sci Transl Med 5: 208ra149

Schludi MH, Becker L, Garrett L, Gendron TF, Zhou Q, Schreiber F, Popper B, Dimou L, Strom TM, Winkelmann J, von Thaden A, Rentzsch K, May S, Michaelsen M, Schwenk BM, Tan J, Schoser B, Dieterich M, Petrucelli L, Holter SM et al. (2017) Spinal poly-GA inclusions in a C9orf72 mouse model trigger motor deficits and inflammation without neuron loss. Acta Neuropathol

Scotti MM, Swanson MS (2016) RNA mis-splicing in disease. Nat Rev Genet 17: 19–32

Shao Q, Liang C, Chang Q, Zhang W, Yang M, Chen JF (2019) C9orf72 deficiency promotes motor deficits of a C9ALS/FTD mouse model in a dose-dependent manner. Acta Neuropathol Commun 7: 32

Shi KY, Mori E, Nizami ZF, Lin Y, Kato M, Xiang S, Wu LC, Ding M, Yu Y, Gall JG, McKnight SL (2017) Toxic PRn poly-dipeptides encoded by the C9orf72 repeat expansion block nuclear import and export. Proc Natl Acad Sci U S A 114: E1111–E1117

Stewart H, Rutherford NJ, Briemberg H, Krieger C, Cashman N, Fabros M, Baker M, Fok A, DeJesus-Hernandez M, Eisen A, Rademakers R, Mackenzie IR (2012) Clinical and pathological features of amyotrophic lateral sclerosis caused by mutation in the C9ORF72 gene on chromosome 9p. Acta Neuropathol 123: 409–17

Tada M, Doi H, Koyano S, Kubota S, Fukai R, Hashiguchi S, Hayashi N, Kawamoto Y, Kunii M, Tanaka K, Takahashi K, Ogawa Y, Iwata R, Yamanaka S, Takeuchi H, Tanaka F (2018) Matrin 3 Is a Component of Neuronal Cytoplasmic Inclusions of Motor Neurons in Sporadic Amyotrophic Lateral Sclerosis. Am J Pathol 188: 507–514

Taylor JP, Brown RH, Jr., Cleveland DW (2016) Decoding ALS: from genes to mechanism. Nature 539: 197–206

van Blitterswijk M, Baker MC, DeJesus-Hernandez M, Ghidoni R, Benussi L, Finger E, Hsiung GY, Kelley BJ, Murray ME, Rutherford NJ, Brown PE, Ravenscroft T, Mullen B, Ash PE, Bieniek KF, Hatanpaa KJ, Karydas A, Wood EM, Coppola G, Bigio EH et al. (2013) C9ORF72 repeat expansions in cases with previously identified pathogenic mutations. Neurology 81: 1332–41

van der Zee J, Gijselinck I, Dillen L, Van Langenhove T, Theuns J, Engelborghs S, Philtjens S, Vandenbulcke M, Sleegers K, Sieben A, Baumer V, Maes G, Corsmit E, Borroni B, Padovani A, Archetti S, Perneczky R, Diehl-Schmid J, de Mendonca A, Miltenberger-Miltenyi G et al. (2013) A pan-European study of the C9orf72 repeat associated with FTLD: geographic prevalence, genomic instability, and intermediate repeats. Hum Mutat 34: 363–73

Van Langenhove T, van der Zee J, Gijselinck I, Engelborghs S, Vandenberghe R, Vandenbulcke M, De Bleecker J, Sieben A, Versijpt J, Ivanoiu A, Deryck O, Willems C, Dillen L, Philtjens S, Maes G, Baumer V, Van Den Broeck M, Mattheijssens M, Peeters K, Martin JJ et al. (2013) Distinct clinical characteristics of C9orf72 expansion carriers compared with GRN, MAPT, and nonmutation carriers in a Flanders-Belgian FTLD cohort. JAMA Neurol 70: 365–73

Van Mossevelde S, van der Zee J, Cruts M, Van Broeckhoven C (2017a) Relationship between C9orf72 repeat size and clinical phenotype. Curr Opin Genet Dev 44: 117–124

Van Mossevelde S, van der Zee J, Gijselinck I, Sleegers K, De Bleecker J, Sieben A, Vandenberghe R, Van Langenhove T, Baets J, Deryck O, Santens P, Ivanoiu A, Willems C, Baumer V, Van den Broeck M, Peeters K, Mattheijssens M, De Jonghe P, Cras P, Martin JJ et al. (2017b) Clinical Evidence of Disease Anticipation in Families Segregating a C9orf72 Repeat Expansion. JAMA Neurol 74: 445–452

Waite AJ, Baumer D, East S, Neal J, Morris HR, Ansorge O, Blake DJ (2014) Reduced C9orf72 protein levels in frontal cortex of amyotrophic lateral sclerosis and frontotemporal degeneration brain with the C9ORF72 hexanucleotide repeat expansion. Neurobiol Aging 35: 1779 e5–1779 e13

Wils H, Kleinberger G, Janssens J, Pereson S, Joris G, Cuijt I, Smits V, Ceuterick-de Groote C, Van Broeckhoven C, Kumar-Singh S (2010) TDP-43 transgenic mice develop spastic paralysis and neuronal inclusions characteristic of ALS and frontotemporal lobar degeneration. Proc Natl Acad Sci U S A 107: 3858–63

Wooley CM, Sher RB, Kale A, Frankel WN, Cox GA, Seburn KL (2005) Gait analysis detects early changes in transgenic SOD1(G93A) mice. Muscle Nerve 32: 43–50

Wright GEB, Collins JA, Kay C, McDonald C, Dolzhenko E, Xia QW, Becanovic K, Drogemoller BI, Semaka A, Nguyen CM, Trost B, Richards F, Bijlsma EK, Squitieri F, Ross CJD, Scherer SW, Eberle MA, Yuen RKC, Hayden MR (2019) Length of Uninterrupted CAG, Independent of Polyglutamine Size, Results in Increased Somatic Instability, Hastening Onset of Huntington Disease. American Journal of Human Genetics 104: 1116–1126

Xiao S, MacNair L, McGoldrick P, McKeever PM, McLean JR, Zhang M, Keith J, Zinman L, Rogaeva E, Robertson J (2015) Isoform-specific antibodies reveal distinct subcellular localizations of C9orf72 in amyotrophic lateral sclerosis. Ann Neurol 78: 568–83

Xu Z, Poidevin M, Li X, Li Y, Shu L, Nelson DL, Li H, Hales CM, Gearing M, Wingo TS, Jin P (2013) Expanded GGGGCC repeat RNA associated with amyotrophic lateral sclerosis and frontotemporal dementia causes neurodegeneration. Proc Natl Acad Sci U S A 110: 7778–83

Yin S, Lopez-Gonzalez R, Kunz RC, Gangopadhyay J, Borufka C, Gygi SP, Gao FB, Reed R (2017a) Evidence that C9ORF72 Dipeptide Repeat Proteins Associate with U2 snRNP to Cause Mis-splicing in ALS/FTD Patients. Cell Rep 19: 2244–2256

Yin SY, Lopez-Gonzalez R, Kunz RC, Gangopadhyay J, Borufka C, Gygi SP, Gao FB, Reed R (2017b) Evidence that C9ORF72 Dipeptide Repeat Proteins Associate with U2 snRNP to Cause Mis-splicing in ALS/FTD Patients. Cell Reports 19: 2244–2256

Zhang K, Donnelly CJ, Haeusler AR, Grima JC, Machamer JB, Steinwald P, Daley EL, Miller SJ, Cunningham KM, Vidensky S, Gupta S, Thomas MA, Hong I, Chiu SL, Huganir RL, Ostrow LW, Matunis MJ, Wang J, Sattler R, Lloyd TE et al. (2015) The C9orf72 repeat expansion disrupts nucleocytoplasmic transport. Nature 525: 56–61

Zhang Y, Chen K, Sloan SA, Bennett ML, Scholze AR, O’Keeffe S, Phatnani HP, Guarnieri P, Caneda C, Ruderisch N, Deng S, Liddelow SA, Zhang C, Daneman R, Maniatis T, Barres BA, Wu JQ (2014) An RNA-sequencing transcriptome and splicing database of glia, neurons, and vascular cells of the cerebral cortex. J Neurosci 34: 11929–47

Zhang YJ, Gendron TF, Ebbert MTW, O’Raw AD, Yue M, Jansen-West K, Zhang X, Prudencio M, Chew J, Cook CN, Daughrity LM, Tong J, Song Y, Pickles SR, Castanedes-Casey M, Kurti A, Rademakers R, Oskarsson B, Dickson DW, Hu W et al. (2018) Poly(GR) impairs protein translation and stress granule dynamics in C9orf72-associated frontotemporal dementia and amyotrophic lateral sclerosis. Nat Med 24: 1136–1142

Zhang YJ, Guo L, Gonzales PK, Gendron TF, Wu YW, Jansen-West K, O’Raw AD, Pickles SR, Prudencio M, Carlomagno Y, Gachechiladze MA, Ludwig C, Tian RL, Chew J, DeTure M, Lin WL, Tong JM, Daughrity LM, Yue M, Song YP et al. (2019) Heterochromatin anomalies and double-stranded RNA accumulation underlie C9orf72 poly(PR) toxicity. Science 363: 707–+

Zu T, Liu Y, Banez-Coronel M, Reid T, Pletnikova O, Lewis J, Miller TM, Harms MB, Falchook AE, Subramony SH, Ostrow LW, Rothstein JD, Troncoso JC, Ranum LP (2013) RAN proteins and RNA foci from antisense transcripts in C9ORF72 ALS and frontotemporal dementia. Proc Natl Acad Sci U S A 110: E4968–77

