## Supplementary Materials and Figures for "Repeat length increases disease penetrance and severity in *C9orf72* ALS/FTD BAC transgenic mice"

*Corresponding author

**Contact information of corresponding author**

### Materials and methods

#### Mate Pair Preparation and Whole Genome Sequencing

Input DNA was received and quantified using a fluorometric-based method specific for double-stranded DNA. The quality of the genomic DNA was evaluated using capillary electrophoresis-based technology (Fragment Analyzer, AATI). Four micrograms of good quality DNA were processed into Illumina-compatible libraries using Nextera Mate Pair Sample Preparation Kit (Illumina) following the Gel-Plus procedure with the following modifications. The fragmented DNA was size-selected for a target range of 5-8 kb fragments using an automated DNA size-selection technology, BluePippin (Sage Science) and the adapter-ligated mate pair fragments were enriched using 10 amplification cycles. Final libraries were quantified using the KAPA Library Quantification Kit (KAPA Biosystem), Qubit Fluorometer (Life Technologies) and 2100 BioAnalyzer (Agilent). Libraries were sequenced on an Illumina HiSeq2500 sequencer using 2 x 125bp cycles.

#### Bioinformatics Analysis for Transgene Integration

All read data was processed prior to alignment. Processing included: 1) ab initio duplicate removal using in-house scripts; 2) adapter identification/clipping and low-quality end trimming (Q=20) using Cutadapt 1.8.1 (Martin, 2011) 3) phiX decontamination by mapping with GEM mapper (Marco-Sola, Sammeth et al., 2012) against the phiX reference (acc. NC_001422.1). Read pairs were further screened for the presence of the Nextera ligation adapter, which was also trimmed. After determination of the library insert size and mate orientation by mapping a subset of the reads against the mouse reference genome (mm10) with the GEM mapper, we decided to consider for downstream analysis the entire set of processed mate-pairs, given that the overall percentage of reads mapping in forward-reverse orientation (representing paired-end contamination) was low (~1%). Processed read data was then mapped, using BWA-MEM(Li, 2013), against a combined index that included, apart from the full mouse reference genome (mm10), the human *C9orf72* sequence and the pCC1 vector used in transfection. Despite the fact that most duplicate sequences have been removed in the initial processing of the data, additional duplicates were identified and marked by the MarkDuplicates tool from Picard (Picard) (from Broad Institute). Integration sites were identified by manual inspection of discordant read-pairs in which one end mapped to the mouse reference and the other end to the *C9orf72* or pCC1 sequences. Analysis of soft-clipped reads at the integration sites allowed determining, with base-pair resolution, the exact breakpoint location in both the mouse genome and the *C9orf72* sequence. Further analysis on structure of the integration was based on read-pairing information and in copy-number analysis of the *C9orf72* and pCC1 sequences. For the latter, both the *C9orf72* and the pCC1 sequences were segmented based on integration breakpoints and deletion site boundaries and, for each segment, average depth of coverage was computed using Depth of coverage from the Genome Analysis Toolkit (GATK) (McKenna A, 2010) The obtained values were then compared to the average depth of coverage over the mouse reference, estimated from a set of intervals overlapping genic regions. For calling variants over both the *C9orf72* and pCC1 sequences, alignments from all four samples were combined into a single BAM file and GATK's HaplotypeCaller was used to produce a gVCF file. Variants were then emitted by GATK's GenotypeGVCFs.

**Transcriptome Sequencing and Data Analysis**

Animals were perfused transcardially with sterile 1X PBS and the frontal cortex was harvested for RNA isolation. RNA was isolated using the Direct-Zol RNA miniprep kit (Zymo Research). RNA-seq libraries were prepared from total RNA (500–700 ng) using the directional RNA-seq kit with NEB Ultra Ribominus (New England Labs) following the manufacturer's protocol. The RNA was then sequenced using the Illumina NextSeq500 machine at the Center for NeuroGenetics, University of Florida.

**RNA Sequencing Data Analysis**

Reads obtained after sequencing were aligned using STAR (Dobin et al., 2013) to either the human genome (hg19) or mouse genome (mm10). Quality analysis of the reads was performed using RSeQC. We used Kallisto (Bray et al., 2016) to obtain transcript per million (TPM) values for some analyses. GENCODE was used for gene annotations from STAR alignment wand RefSeq was used for Kaliisto gene annotations. DEseq2 was used to measure gene expression differences, run using the Bioconductor package for R. We used MISO based on MISO annotations version 2.0 to identify changes in alternative splicing (Katz et al., 2010). Only events with Bayes factor ≥5 for at least one pair-wise comparison were considered. Events were filtered with $|\Delta\Psi| \geq 0.10$ compared to the NT mice and the venn diagrams were plotted using Biovenn. To identify enriched biological pathways, gene ontology analysis was performed using Metascape using gene sets from GO biological processes and was confirmed using DAVID annotation using default statistical thresholds (P-value<0.01, maximum enrichment >1.5). Graphs were plotted on R using ggplot2. C9-ALS patient data was obtained from Prudencio et al. Data from frontal cortex on autopsy tissue samples were filtered based on RIN scores and samples used were: C9(+)-ALS - SRR1927020, SRR1927022, SRR1927024, SRR1927026, SRR1927028, SRR1927030, SRR1927032 and SRR1927034; Control - SRR1927056, SRR1927058, SRR1927060, SRR1927062, SRR1927064, SRR1927066, SRR1927068, SRR1927070, SRR1927071

**Cell Type Enrichment Analysis**

The index for relative expression of genes in different cell types was set up using publicly available data on cell type enrichment in the mouse cortex. Since the differences between the oligodendrocyte cell types were subtle, we combined myelinating oligodendrocytes, oligodendrocyte precursor cells and mature oligodendrocytes into one classification "oligodendrocyte". For a gene to belong to a particular cell type, expression had to be >20FPKM and logFC between any sample and mean of all other samples > 2. Bayesian statistical model was used to calculate the proportions of cell types within the current dataset. The python package Pymc3 was used to perform these analyses and Matplotlib was used to generate the graphs.

**Motif Analysis**

To analyze the number of enriched motifs, we used the monotonically changing cassette exons from alternatively spliced events generated from MISO with a Bayes factor>5 and $|\Delta\Psi| \geq 0.10$ in NT and Acute animals. Fisher's exact test and Bonferroni correction were performed and the p adjusted<0.05 was set as the significance threshold. The regions 250 bp upstream, exon, 250 bp downstream was used to count for all possible combination of 4-mers. We used Fisher's exact test to identify motifs that are enriched on depleted specifically in Acute mice when compared to NT mice.

**Splicing analysis**

For splicing analyses, mm10 annotated events were run on MISO to identify alternatively spliced events to identify skipped exon (SE) events. Delta PSI values for NT vs C9 mean PSIs and NT vs Acute mean PSIs were calculated and reported in Table 2. Wilcoxon rank-sum test was also run for both comparisons. Delta PSI cutoff was set at 0.1 (or -0.1) and a rank sum p value < 0.05 to establish significance. GO analysis was run with metascape.org.

**Southern blot**

Tail genomic DNA (gDNA) extraction and Southern blot experiments were performed following previously published protocols in Liu et al., 2016. Briefly, 10μg of gDNA extracted from the tail or brain was digested with EcoRI and BamHI overnight at 37°C. The digested gDNA samples were then run on a 0.7% agarose gel for ~5-6hours, depurinated (0.2N HCl), denatured (1.5M NaCl, 0.5M NaOH) and neutralized (1.5M NaCl, 0.5M Tris HCl) for 15 mins each. DNA was transferred overnight by capillary blotting to a positively charged nylon membrane and was UV cross-linked the next morning. For hybridization, the membrane was prehybridized for 1 hour using Amersham Rapid-Hyb buffer (GE Healthcare). The probe was labeled using dCTP-$P^{32}$ using the random primed DNA labeling kit (Invitrogen) and was hybridized to the membrane. After 3 hours of hybridization, the membrane was washed with 2x SSC, 0.1%SDS for 20 minutes at room temperatures and then with 0.2x SSC, 0.1%SDS solution at 65°C two times for 15 minutes each. Radioactivity was visualized on an X-ray film after 2-3 days of exposure at -80°C.

**Quantitative real time PCR**

Total RNA was isolated from mouse frontal cortex and spinal cord tissues with TRIzol (Invitrogen). Following DNase treatment (Ambion) using manufacturer's protocol, cDNA was prepared using the SuperScript III RT kit (Invitrogen) and random-hexamer primers (Applied Biosystems). Two-step quantitative RT-PCR was performed on a MyCycler Thermal Cycler system (Bio-Rad) using SYBR Green PCR Master Mix (Bio-Rad). See primer lists in Table 4.

**Mouse generation and maintenance**

C9-BAC transgenic mice were mated with FVB mice obtained from Jackson Laboratory. Pups were genotyped based on genomic DNA extracted from the tail using the primers C9-GT F and R (Table 3) and the protocol previously described by Liu et al., 2016.  The repeat size of each mouse in the study was determined using Southern blotting described above. Behavioral tests were performed at 12, 16 and 24 weeks of age. DigiGait and open field analyses were performed based on the manufacturer's protocol. For molecular and pathological staining, mice were perfused transcardially using 1x PBS and tissues were embedded in 10% formalin or OCT frozen in cold 2-methylbutane for further analyses.

**Fluorescent In Situ Hybridization (FISH)**

Ten μm frozen sections embedded in OCT were cut on the cryostat. Frozen sections were fixed in 4% PFA in PBS for 20 min and incubated in prechilled 70% ethanol for 30 min or longer at 4°C. Following rehydration in 40% formamide in 2× SSC at room temperature for 10 min, the slides were prehybridized with hybridization solution (40% formamide, 2× SSC, 20 μg/mL BSA, 100 mg/mL dextran sulfate, 250 μg/mL yeast tRNA, 2 mM Vanadyl Sulfate Ribonucleosides) for 30 min at 55 °C and then incubated with 200 ng/mL of denatured DNA probe ($(C2G4)_3$-Cy3 for sense foci and $(G4C2)_3$-Cy3 for antisense foci) in hybridization solution at 55 °C. After 3h of hybridization, the slides were washed three times with 40% formaminde in 2× SSC and briefly washed one time in PBS. Autofluorescence of lipofuscin was quenched by 0.25% of Sudan Black B in 70% ethanol. Slides were mounted with ProLong mounting medium containing DAPI (Invitrogen) and imaged on LSM880 Confocal microscope.

**Immunohistochemistry**

Five μm sections were deparaffinized in xylene and rehydrated through gradient ethanol solution. Antigen retrieval was performed by incubating the slides in a steamer with 10mM citrate buffer (pH 6.0) for 30 min or with 10mM EDTA (pH 6.4) for 12 min. The slides were cooled down to RT and then washed for 10 min in running tap water. The slides were incubated in 95-100% formic acid for 5 min and subsequently washed for 10 min in running tap water. To eliminate nonspecific binding and excessive background, slides were blocked with a serum free block or rodent block (Biocare Medical) for 15 min. Primary antibody diluted in 1:10 blocking solution was applied on the slides and incubated overnight at 4°C (see below for dilution information). Slides were washed three times with 1XPBS and incubated with linking reagent (streptavidin or alkaline phosphatase; Covance) or biotinylated rabbit anti-goat IgG (Vector Labs) for 30 min at room temperature. After washing with 1XPBS, these sections were then incubated in 3% hydrogen peroxide (in methanol) for 15 min to eliminate any endogenous peroxidase activity. After washing in running tap water for 15 min, labeling reagent (HRP, Covance; Vectastain ABC-AP kit) was then applied to the slides for 30 min at room temperature. The slides were developed with NovaRed or DAB (Vector Labs) and then were counterstained with hematoxylin (modified Harris, Sigma Aldrich), rehydrated in gradient ethanol solution and coverslipped for visualization. Images were taken on the Olympus BX51 microscope using the cellSense software. For hematoxylin and eosin staining, the slides were deparaffinized in xylene and rehydrated through gradient ethanol solution. The slides were then soaked in hematoxylin (modified Harris, Sigma Aldrich) for 1 min and washed in running distilled water for 10 min. Next, the slides were immersed in Eosin Y (Sigma Aldrich) for 30 sec and washed in distilled water for 10 min. The slides were dehydrated and coverslipped before visualization. For cresyl violet staining, slides were deparaffinized in xylene and subsequently rehydrated in gradient ethanol solution. The slides were incubated in 0.25% cresyl violet at

60°C for 8-10 min and differentiated in 95% ethanol for 1-5 mins. Slides were then immersed in 100% ethanol and xylene and coverslipped for visualization.

### Immunofluorescence

Frozen sections were warmed up at room temperature for 2 hours. Slides were fixed with 4% PFA at room temperature for 10 minutes and immediately permeabilized with ice-cold 1:1 methanol-acetone for 5 minutes at -20°C. Slides were blocked with background sniper at RT for 30 min to 1 h and incubated overnight with primary antibody (polyGA 1:2000, polyGP 1:1000, generously donated by Neurimmune Inc.) prepared in a 1:10 dilution of background sniper. The slides were washed three times with 1xPBS the next day and incubated with secondary antibody (Cy3-anti-human IgG antibody, 1:5000) for 1-2 hours. The slides were washed with 1x PBS and mounted with Prolong containing DAPI. Slides were imaged with a Zeiss LSM880 Confocal microscope.

#### Western Blot

Brain lysates were solubilized with RIPA buffer (1% sodium deoxycholate, 1% Triton X-100, 50mM Tris pH 7.5, 150mM NaCl and proteinase inhibitor). 10μl of lysates were run on a 4-12% Bis-Tris gel and transferred to a nitrocellulose membrane. The membrane was blocked in 5% milk diluted in PBST (1XPBS with 0.05% Tween-20). The membrane was then incubated with the primary antibody (α-C9orf72, 1:1000) overnight at 4°C. The blots were washed three times with 1XPBS the next morning and were incubated with secondary antibody for 1h at RT. The membrane was developed using ECL prime and was visualized for signal.

### Image J quantification

Quantification was performed by two independent blinded investigators using the cell counter plugin in ImageJ (National Institute of Health). Serial sections 20-30μm apart were stained, imaged and quantified for subsequent analyses.

**Behavioral analyses**

DigiGait, open field analyses and scoring criteria for cage behavioral assessments were performed as previously described in Liu et al., 2016.

**Statistics**

GraphPad Prism 7 was used to perform the statistical analyses in the manuscript. Significance threshold was set at p<0.05. The significance values were set as ns=not significant p>0.05, p<0.05 – "*", p<0.01 – "**", p<0.001 – "***", p<0.0001 – "****". Comparisons between groups was performed using mean±SEM values using one-way ANOVA and multiple comparisons. Significance in survival was measured using the Log Mantel-Cox test. DigiGait analyses were performed using multiple comparisons.

### Supplementary figures

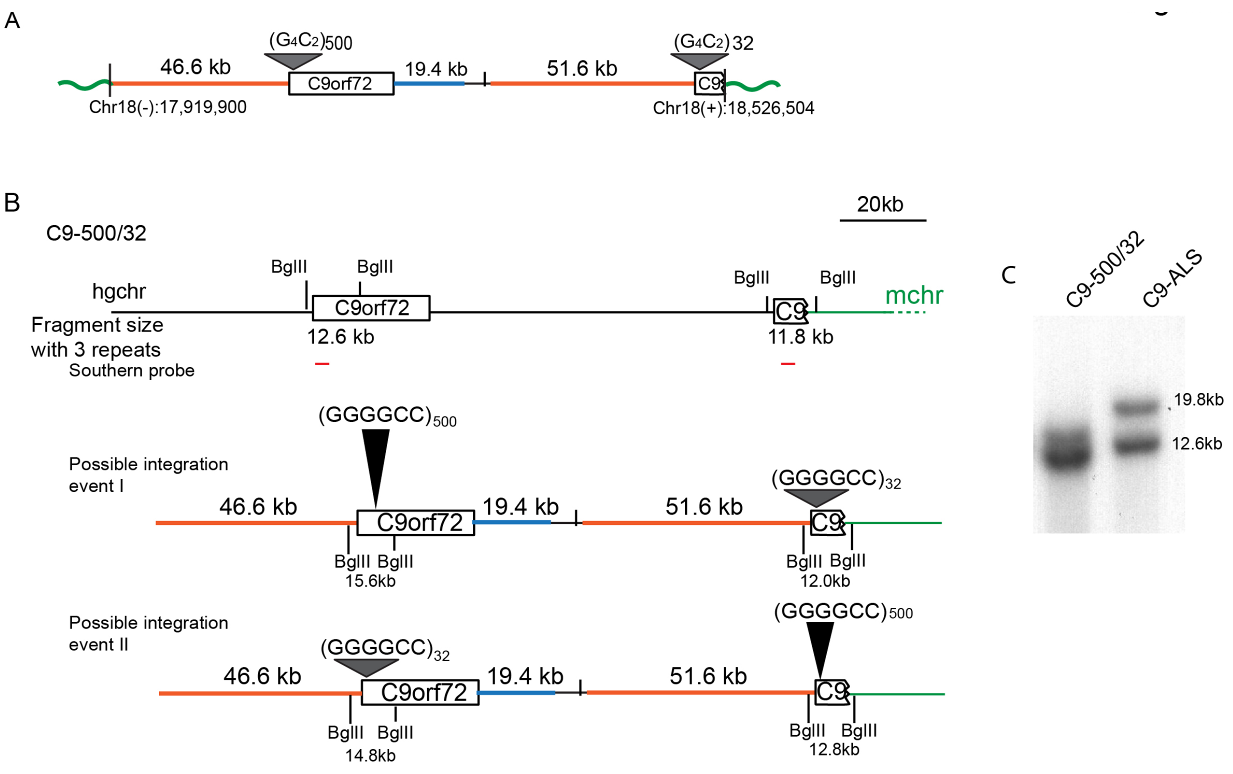


**Figure S1: Analysis of transgene integration sites in C9-500/32 line.**

A) Schematic diagram of transgene integration in C9-500/32 line.

B) Schematic diagrams indicating the possible location of GGGGCC repeats with 500 or 32 copies.

A, B) Orange and blue lines represent human transgene sequence upstream and downstream of the expansion mutation, respectively. Green lines denote mouse chromosomal regions.

C) Southern blot of genomic DNA digested with BglII shows that the larger repeat was integrated into the full-length transgene as predicted in possible integration event I.

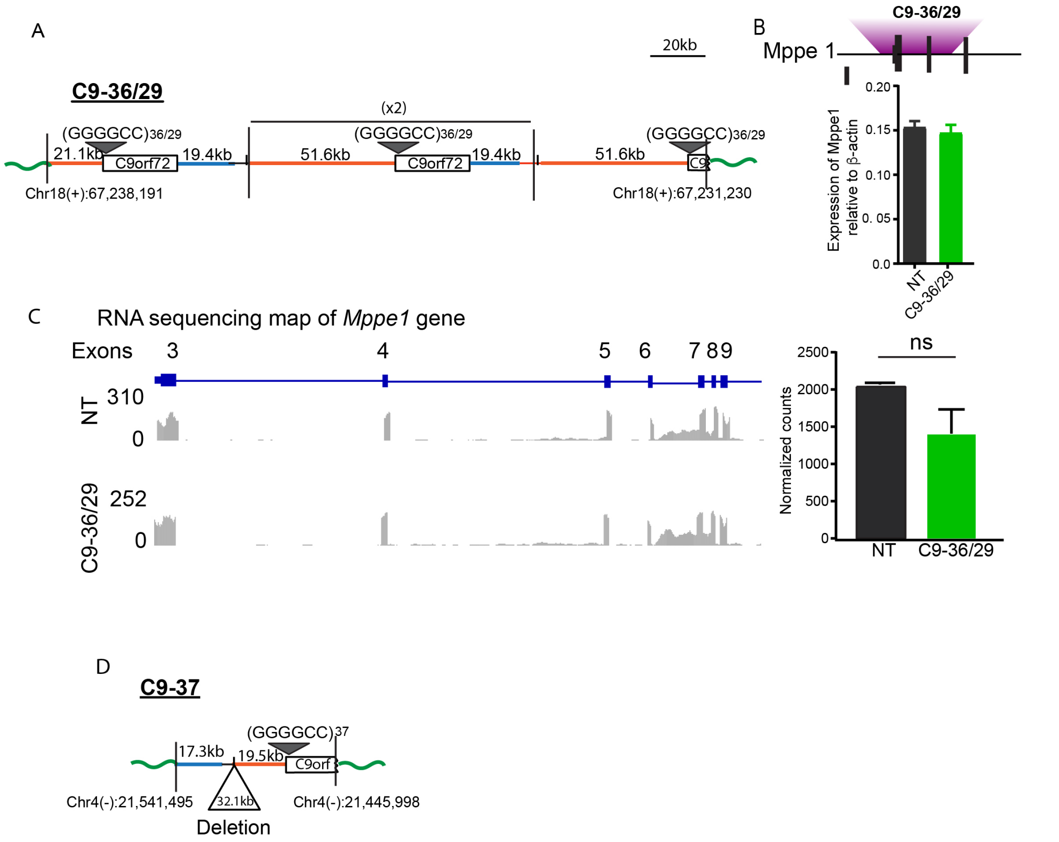


**Figure S2: Transgene integration in C9-36/29 and C9-37 lines.**

A) Schematic diagram showing transgene integration site in the C9-36/29 line contains four copies of BAC transgene integrated in *Mppe1*.

B) Schematic diagram showing location of transgene within *Mppe1*, qPCR shows no change in expression of *Mppe1* gene upon transgene integration.

C) RNA sequencing of C9-36/29 mice showed no change in coverage over *Mppe1* gene.

D) Schematic diagram showing transgene integration site in C9-37 mice.

A, D) Orange and blue lines represent human transgene sequence upstream and downstream of the expansion mutation, respectively. Green lines denote mouse chromosomal regions.

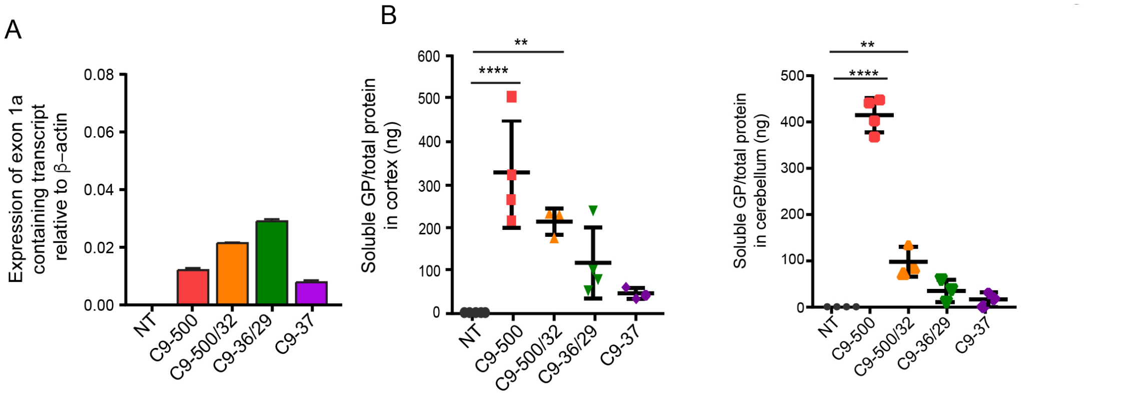


**Figure S3: RNA and RAN protein levels in C9-BAC transgenic mice.**

A) RT-qPCR shows levels of exon1a containing transcripts in C9-BAC transgenic lines.

B) MSD immunoassay to measure levels of soluble GP in cortical brain lysates.

C) MSD immunoassay to measure levels of soluble GP in cerebellar brain lysates.

Data information: Statistical analyses for (A), (B) and (C) was done using a one-way ANOVA with multiple comparison, Bonferroni correction, Mean ± SEM, *p<0.05, **p<0.01, ****p <0.0001.

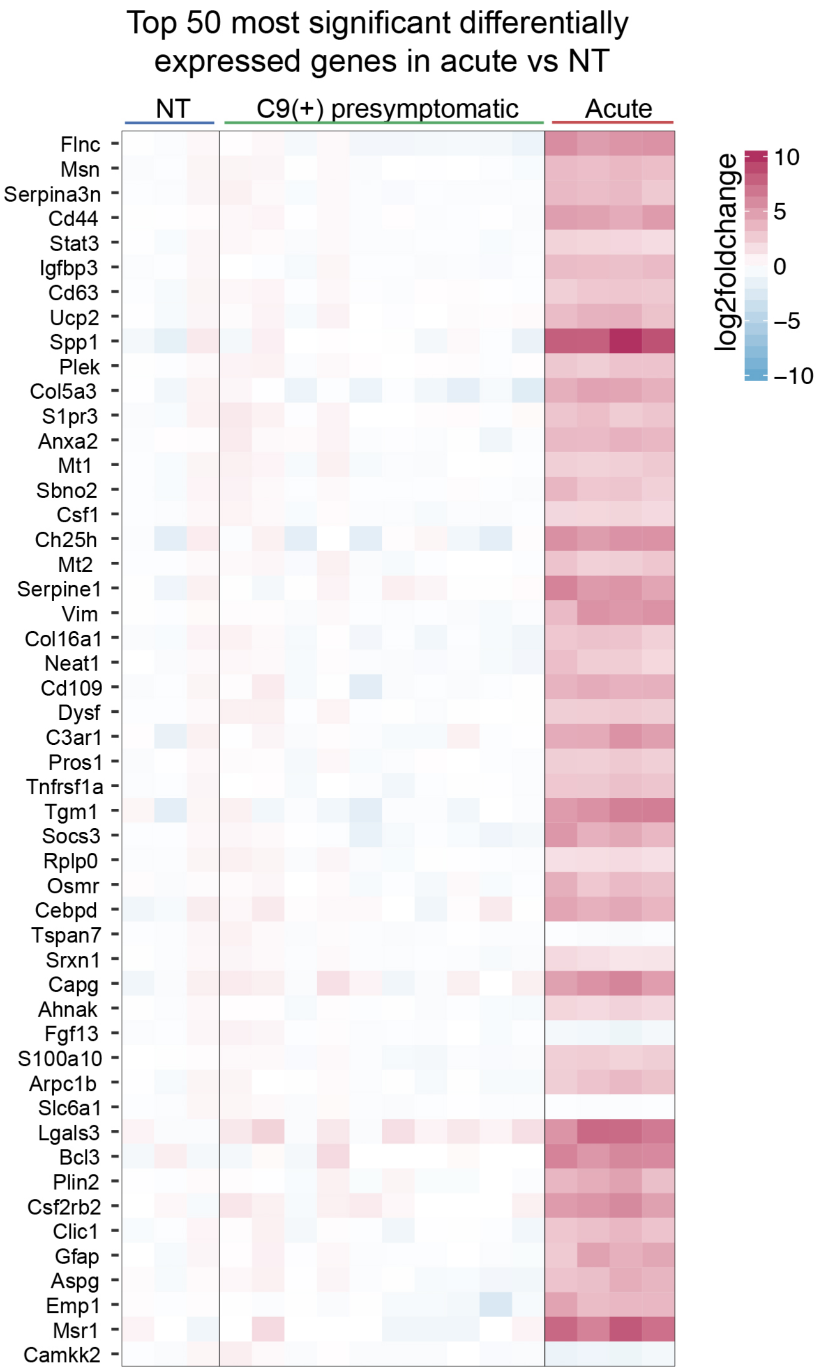


**Figure S4: Transcriptome changes in Acute C9-BAC mice**.

Heat map shows the top 50 differentially expressed genes in Acute vs NT mice and their relative expression in NT and C9(+) presymptomatic mice.

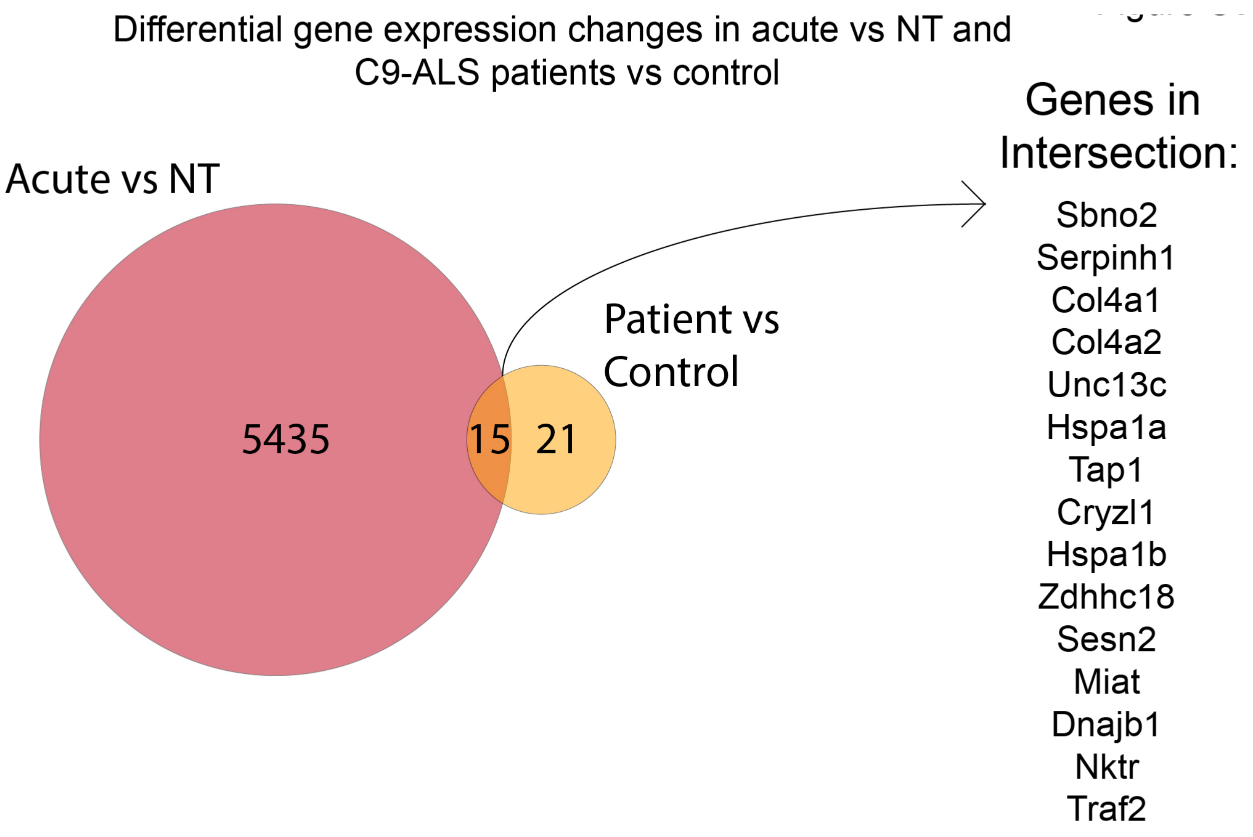


**Figure S5: Differential gene expression changes in Acute vs NT mice and C9-ALS patients' vs controls.**

Venn diagram showing overlap between differential gene expression measured from Acute vs NT mice and C9-ALS patients' vs controls. The genes from the intersection category are listed.

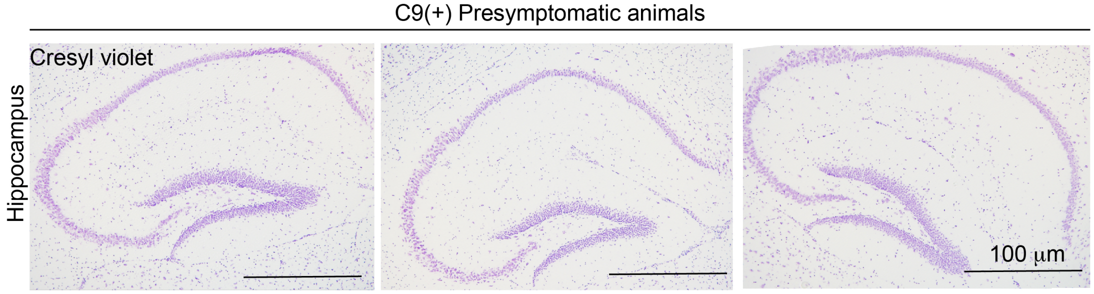


**Figure S6: Cresyl violet staining of the hippocampus in C9(+) presymptomatic mice.**

No overt neuropathology was observed in the hippocampus of C9(+) presymptomatic animals.

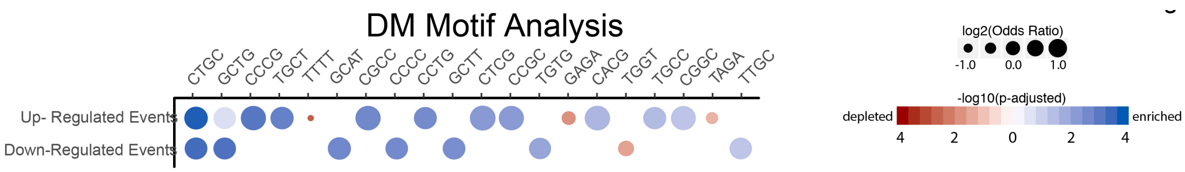


**Figure S7: Motif analysis from alternative splicing dataset obtained from TA muscle of DM1 patients.**

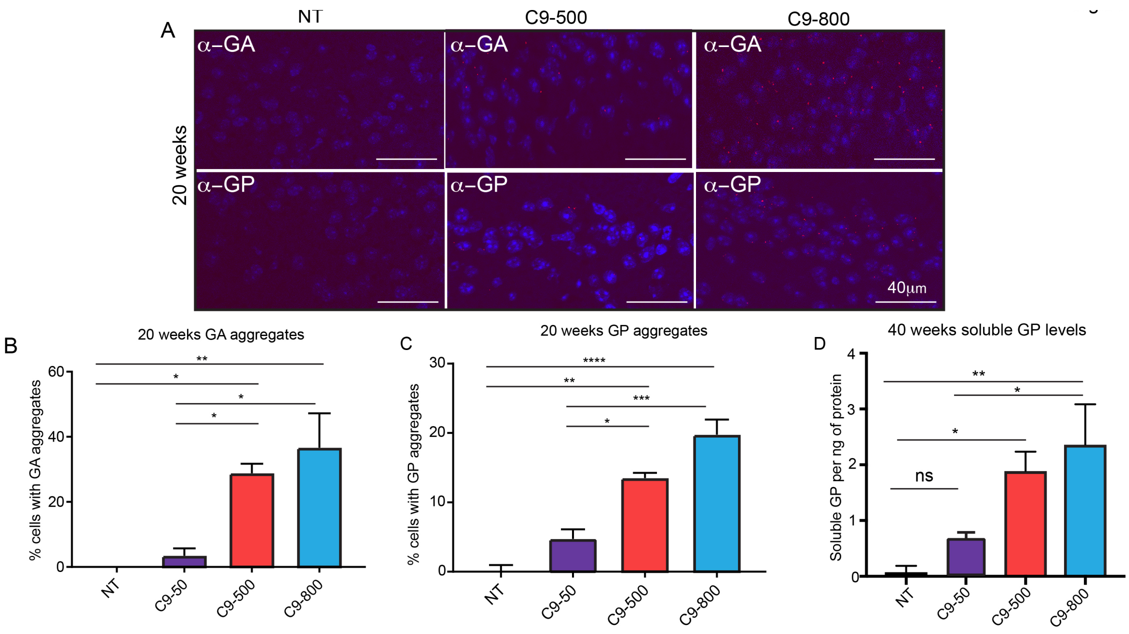


**Figure S8: Molecular changes in allelic series sublines.**

A) Representative images of GA and GP aggregates in the retrosplenial cortex of C9 BAC mice.

B, C) Quantification of GA and GP aggregates measured in the retrosplenial cortex at 20 weeks of age.

D) MSD immunoassay to measure levels of soluble GP in brain lysates at 40 weeks in allelic series sublines.

Data information: Statistical analyses for (B), (C) and (D) were done using a one-way ANOVA with multiple comparison, Bonferroni correction, Mean ± SEM, $^*p<0.05$, $^{**}p<0.01$, $^{****}p<0.0001$.
